# A Single-Cell Tumor Immune Atlas for Precision Oncology

**DOI:** 10.1101/2020.10.26.354829

**Authors:** Paula Nieto, Marc Elosua-Bayes, Juan L. Trincado, Domenica Marchese, Ramon Massoni-Badosa, Maria Salvany, Ana Henriques, Elisabetta Mereu, Catia Moutinho, Sara Ruiz, Patricia Lorden, Vanessa T. Chin, Dominik Kaczorowski, Chia-Ling Chan, Richard Gallagher, Angela Chou, Ester Planas-Rigol, Carlota Rubio-Perez, Ivo Gut, Josep M. Piulats, Joan Seoane, Joseph E. Powell, Eduard Batlle, Holger Heyn

**Affiliations:** CNAG-CRG, Centre for Genomic Regulation (CRG), Barcelona Institute of Science and Technology (BIST), Barcelona, Spain; Institute for Research in Biomedicine (IRB Barcelona), The Barcelona Institute of Science and Technology, Barcelona, Spain; Garvan-Weizmann Centre for Cellular Genomics, Garvan Institute of Medical Research, Sydney, Australia; St Vincent’s Hospital Clinical School, University of New South Wales, Sydney; St Vincent’s Hospital Sydney, Darlinghurst, Australia; St Vincent’s Hospital Sydney, Darlinghurst, Australia; University of Notre Dame, Sydney, Australia; Cancer Diagnosis and Pathology Group, Kolling Institute of Medical Research, Royal North Shore Hospital, St Leonards NSW 065 Australia; NSW Health Pathology, Department of Anatomical Pathology, Royal North Shore Hospital, Sydney NSW 2065 Australia; University of Sydney, Sydney, NSW 2006 Australia; Vall d’Hebron Institute of Oncology (VHIO), Vall d’Hebron University Hospital, 08035 Barcelona, Spain; Medical Oncology Department, Institut Català d’Oncologia - ICO; Clinical Research in Solid Tumors Group - CREST, Bellvitge Biomedical Research Institute IDIBELL-OncoBell; CIBERONC; L’Hospitalet de Llobregat, Barcelona, Spain; Institució Catalana de Recerca i Estudis Avançats (ICREA), Barcelona, Spain; Universitat Autònoma de Barcelona (UAB), Barcelona, Spain; CIBERONC, Barcelona, Spain; UNSW Cellular Genomics Futures Institute, University of New South Wales, Sydney, Australia; Universitat Pompeu Fabra (UPF), Barcelona, Spain

**Keywords:** Cancer, Immuno-oncology, Tumor microenvironment, Single-cell genomics, Transcriptomics, RNA sequencing, Spatial transcriptomics, Immuno-therapy, Digital pathology

## Abstract

The tumor immune microenvironment is a main contributor to cancer progression and a promising therapeutic target for oncology. However, immune microenvironments vary profoundly between patients and biomarkers for prognosis and treatment response lack precision. A comprehensive compendium of tumor immune cells is required to pinpoint predictive cellular states and their spatial localization. We generated a single-cell tumor immune atlas, jointly analyzing >500,000 cells from 217 patients and 13 cancer types, providing the basis for a patient stratification based on immune cell compositions. Projecting immune cells from external tumors onto the atlas facilitated an automated cell annotation system for a harmonized interpretation. To enable *in situ* mapping of immune populations for digital pathology, we applied *SPOTlight*, combining single-cell and spatial transcriptomics data and identifying striking spatial immune cell patterns in tumor sections. We expect the tumor immune cell atlas, together with our versatile toolbox for precision oncology, to advance currently applied stratification approaches for prognosis and immuno-therapy.

## Introduction

Single-cell RNA sequencing (scRNA-seq) techniques are powerful tools for the unbiased charting of cellular phenotypes (Lafzi et al. 2018). Analyzing transcriptome profiles of individual cells enables the fine-grained annotation of cell types and cellular states, as well as charting the composition of complex samples. Interrogating healthy tissues at single-cell resolution provides a reference atlas of normal tissue organization (Regev et al. 2017) and defines variability across individuals (van der Wijst et al. 2018) or during development (Park et al. 2020; Popescu et al. 2019; Asp et al. 2019) and aging (Salzer et al. 2018; Tabula Muris Consortium 2020). Diseased tissues display an additional layer of complexity, presenting cell type composition shifts and newly emerging disease-specific phenotypes (Vieira Braga et al. 2019; Ramachandran et al. 2019; Chua et al. 2020). In cancer, in addition to diverse neoplastic cell states (Tirosh et al. 2016b; Patel et al. 2014), the remodeling of the host tissue microenvironment has been characterized using scRNA-seq (Tirosh et al. 2016a; Puram et al. 2017). Phenotyping single cells from the tumor microenvironment (TME) has led to the identification of cancer-specific stromal cell states and supported their contribution to tumor progression. Functional and integrative analysis further support dependencies of stromal and cancer cells and their predictive value for patient outcome. In particular, cancer-associated fibroblasts (CAFs) (Calon et al. 2015; Merlos-Suárez et al. 2011) and tumor-resident immune cells (Yofe et al. 2020; Fridman et al. 2012) have been identified as biomarkers for patient stratification and, importantly, as an effective target for therapeutic intervention in oncology (Tumeh et al. 2014; Tauriello et al. 2018). Clinically most advanced, immune cells are now target of immuno-therapy (e.g. immune checkpoint inhibitors, ICI), stimulating the immune system to recognize and destroy cancer cells.

Single-cell transcriptomics has critically advanced our understanding of immune cell diversity in the TME by generating high-resolution landscapes of different cancer types. Combined with T-cell receptor (TCR) genotyping and receptor-ligand screening (Mimitou et al. 2019), scRNA-seq identified transient immune cell states and dynamic tissue re-modelling. Seminal studies include the scRNA-seq-based immunophenotyping of breast cancer, describing a tumor-specific heterogeneity and expansion of T-cell states; supporting a continuous cell activation towards terminal exhaustion, rather than discrete cellular states (Azizi et al. 2018). In melanoma, similar dynamics were shown to activate transitional and dysfunctional T-cell states (Li et al. 2019). Moreover, single-cell immuno-phenotyping identified tumor-specific T-cell states, which are predictive for ICI-based therapy outcome (Sade-Feldman et al. 2018). A lung cancer study identified an immune activation module characterized through high frequencies of PDCD1+ CXCL13+ activated T cells, IgG+ plasma cells, and SPP1+ macrophages, introducing the concept of an immune cell composition-based patient stratification to complement current genomic (e.g. mutational load) or biomarker (e.g. PDL1 or CTLA4) strategies (Leader et al. 2020).

In more general terms, single-cell sequencing-based immuno-phenotyping identified cancer-specific states and composition biases across all major immune cell types that co-localize with cancer cells (Qian et al. 2020). Major alterations have been described for T-cells (Yost et al. 2019), B-cells (Helmink et al. 2020), tumor-associated macrophages (TAMs) (Lee et al. 2020), and dendritic cells (Lavin et al. 2017), suggesting a global perturbation of the immune system in cancer. Comprehensively understanding the causes and consequences of perturbed immune cell function across cancer types could provide the basis to identify novel therapeutic targets and could lay the ground for an immune-based patient stratification in precision oncology.

However, it remains challenging to generalize findings across cancer types. There are no standardized analysis pipelines and annotation systems, resulting in datasets characterized with varying granularity and nomenclature. Cell annotation is especially challenging when describing novel phenotypes (e.g. cancer-specific cell states), leading to discrepancies in cell labels between studies. Moreover, single-cell studies followed different protocols, resulting in technical biases that challenge meta-analyses (Mereu et al. 2020; Massoni-Badosa et al. 2020). On the other hand, recent technical advances enable the scaling of scRNA-seq experiments to larger cell numbers and patient cohorts (Prakadan et al. 2017), providing a rich resource to identify commonalities and peculiarities of immune cells across cancer types. Joining these data into a reference tumor immune cell atlas is an intriguing window of opportunity for a standardized annotation and an immune cell-based patient stratification. Moreover, emerging research fields, such as digital pathology, could benefit from a universally applicable reference. When combined with integrative computational tools and spatial datasets, such reference can be predictive for the presence and spatial organization of immune cells in tumors.

Current ICI stratification strategies involve the assessment of the tumor mutational burden or inflammation signatures. However, despite having favorable immune profiles many tumors do not respond to treatments, suggesting additional mechanisms that confer resistance to therapy. In this regard, the spatial distribution of immune cells has proven to be important for ICI response, with excluded tumors blocking effective immune cell action, despite the presence of favorable cell types at their boundaries (Chen and Mellman 2017). In contrast, tumors invaded by reactive immune cells that clonally expand have shown increased response rates. However, current spatial immune profiling approaches enable only targeted profiling of mRNAs or proteins. On the other hand, latest spatial transcriptomics (ST) techniques (Ståhl et al. 2016; Rodriques et al. 2019) provide unbiased transcriptome-wide profiles, but average gene expression profiles of multiple cells.

In this work, we generate a consensus atlas of the tumor immune microenvironment through the integration of scRNA-seq datasets from 13 cancer types. We jointly analyzed >500,000 cells from 217 patients, including major (e.g., T-cells, B-cells and macrophages) and minor (e.g. proliferating and dendritic cells) cell types and states infiltrating human tumors. We stratified cell types into subpopulations and annotated a fine-grained map of immune cell states. We observed high cell state heterogeneity across patients and cancer types, enabling a patient stratification based on the immune composition. This immune-based classification identified pan-cancer subtypes exclusively enriched in cytotoxic and terminally exhausted CD8 T-cells, T-helper 17 (Th17), recently activated CD4 T-cells, M2 tumor-associated macrophages (TAMs) or B-cells and plasma B-cells. We further envision the atlas to serve as a reference for a harmonized annotation of external datasets, which we tested by projecting single cells from primary human tumors and metastases as well as mouse immune-therapy models onto the atlas. To enable *in situ* mapping of atlas populations in tumor sections, we applied *SPOTlight* (Elosua et al. 2020), a computational tool for the integration of reference single-cell and ST datasets. Detected striking regional immune cell enrichments in tumor clones and adjacent areas, we foresee an application of the atlas in digital pathology to guide immune-therapy decisions.

## Results and Discussion

### Generating a tumor immune cell atlas

Single-cell transcriptome profiling of tumors provides an unbiased overview of the heterogeneity of cancer cells and their microenvironment. Following sample dissociation, single cells enter scRNA-seq processes either directly or following the enrichment of specific cell types (e.g. tumor or CD45+ immune cells). To generate a comprehensive tumor immune cell atlas of human cancers, we collected scRNA-seq datasets from 13 different cancer types, 217 patients and 526,261 cells. In detail, we processed data from breast carcinomas (BC) (Azizi et al. 2018), basal cell and squamous cell carcinomas (BCC) (Yost et al. 2019), endometrial adeno-(EA) and renal cell carcinomas (RCC) (Wu et al. 2020), intrahepatic cholangio-(ICC) and hepatocellular carcinomas (HCC) (Ma et al. 2019; Zhang et al. 2019), colorectal cancers (CRC) (Lee et al. 2020; Wu et al. 2020), pancreatic ductal adenocarcinomas (PDAC) (Peng et al. 2019), ovarian cancers (OC) (Schelker et al. 2017), non-small-cell lung cancers (NSCLC) (Lavin et al. 2017; Wu et al. 2020; Lambrechts et al. 2018), and cutaneous (CM) and uveal (UM) melanomas (Li et al. 2019; Sade-Feldman et al. 2018; Durante et al. 2020) (**Supplemental Table 1, Fig. 1A**). For a cell type stratification and consistent annotation, the datasets were analyzed separately before joining immune cells into a pan-cancer tumor immune reference atlas (**Supplemental Fig. 1,2**). In line with previous findings, immune cells clustered by cell identity rather than patient origin, allowing the straightforward subsetting of the immune compartment. We hypothesize that joining cells from different cancer types into a single reference dataset may define commonalities and harmonize annotations between studies.

**Figure 1.**
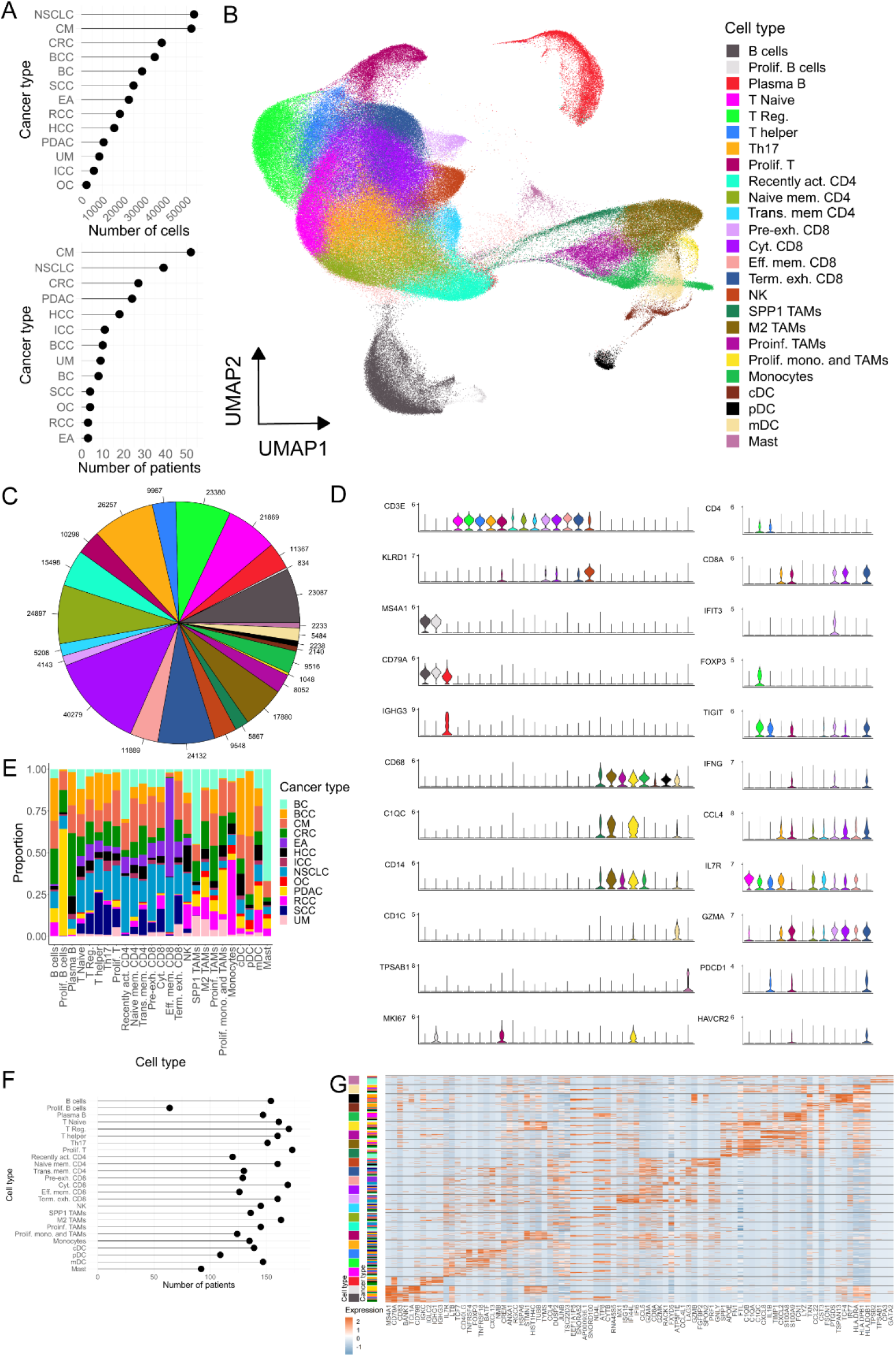
Characterization of the tumor immune cell atlas. **(A)** Number of cells (top) and patients (bottom) per cancer type included in the atlas. **(B)** UMAP of 317,111 immune cells from 13 cancer types colored by annotated cell type. **(C)** Total number of cells of each immune cell type/state; color code as in (A). **(D)** Marker gene expression levels for broader cell types (left) and only T-cells states (right); color code as in (**A**). **(E)** Cancer type proportions for each annotated cell type/state. **(F)** Number of unique patients representing each cell type/state in the atlas. **(G)** Expression of the top 4 differentially expressed genes per cell type/state; colored by cell type (as in A) and cancer type (as in E).

To this end, we integrated 317,111 immune cells using canonical correlation analysis (Butler et al. 2018) (**Fig. 1B**). This approach identifies common cellular phenotypes and allows merging datasets from different studies and technologies for joint analyses. Following integration, cells separated into 25 clusters representing major immune cell types, including 12 T-cell, 5 macrophage/monocyte, 3 dendritic cell (DC), 3 B- and plasma B-cell, 1 natural killer (NK), and 1 Mast cell cluster (**Fig. 1B,C**). To test the robustness of the clusters and their associated signatures, we trained a random forest (RF) classifier to predict cell annotation based on the 25 signatures and performed a 5-fold cross-validation to assess biases and variance. The mean accuracy and kappa statistic across folds were 0.76 (sd = 0.0048) and 0.75 (sd = 0.0050), respectively, a 3.0-fold and 4.2-fold increase with respect to random signatures, and comparable to values obtained in other high-quality atlases (**Supplemental Fig. 3A,B**) (Zeisel et al. 2018). In addition, both the interquartile range (IQR) and the range of accuracies were lower when using cell type-specific signatures, suggesting a low variance across test sets. Finally, stratifying the accuracies by cell type revealed that the low degree of misclassification corresponded to cell types within the same lineage, which share several markers (**Supplemental Fig. 3C**). Taken together this suggests that the clustering of the atlas displays an optimal balance between granularity and robustness.

Clusters were manually annotated using canonical markers and curated gene signatures that defined their identities (**Supplemental Table 2**). Exemplarily, CD4 T-cells representing regulatory, helper and naive states were defined by FOXP3, CXCL13 and SELL marker gene expression, respectively (**Fig. 1D**, **Supplemental Fig. 4**). Cytotoxic CD8 T-cells expressed high levels of GZMA, while their pre- and terminally exhausted subtypes expressed IFIT3 and HAVCR2, respectively. Macrophages split in different subtypes, such as M2, SPP1 and pro-inflammatory states, with C1QB, SPP1 and CXCL8 as respective marker genes. We further observed actively dividing lymphoid and myeloid cell types with MKI67 and STMN1 as common markers for proliferation. Plasma B-cells specifically expressed IGKC, which distinguished them from naive/memory B-cells expressing MS4A1. The large number of immune cells enabled the identification of rare cell states, such as proliferative B-cells (834 cells) and macrophages (1048 cells) as well as plasmacytoid and conventional DC populations (2140 and 2238, **Fig. 1C**). All cell types were present in multiple cancer types and patients (**Fig. 1E,F**), which supports the robustness of the integration approach. Our analyses revealed that several cell states such as naïve, proliferative, central memory and terminally exhausted T-cell subtypes and all macrophage states were abundant all tumors, highlighting common mechanisms that operate in the TME independently of the tissue of origin and mutational background. Yet, we also found cell states enriched in specific cancer types, suggesting cancer-specific immune environments (**Fig. 1E**). In particular, effector memory CD8 T-cells were frequently found in EA and NSCLC, proliferating B-cells were very abundant in PDAC, whereas Mast cells were mainly detected in BC. In summary, we built a catalogued of immune cell types and state markers present in the TME of multiple cancer types (**Fig. 1G** and **Supplemental Table 3**). This resource may help annotate future single-cell tumor datasets.

### Tumor stratification by immune cell composition

Tumors of the same cancer type have been described to be heterogeneous, presenting distinct genetic and epigenetic alterations as well as gene expression signatures, allowing their stratification into subtypes. Cancer subtypes have a clear impact on clinical management being predictive for patient prognosis and therapy response (e.g. consensus molecular subtypes in CRC; CMS) (Guinney et al. 2015). To date, gene expression subtypes are defined through the analysis of bulk tumor samples and are frequently characterized by gene signatures of stromal or immune cell types (Calon et al. 2015). Single-cell resolved tumor maps confirmed this contribution of the TME to the subtype classification and allowed an even more fine-grained interpretation of subtype composition (Lee et al. 2020).

Detecting varying immune cell compositions within cancer types, but conserved profiles across cancers, we sought to establish a pan-cancer immune classification system. We used immune cell type and state frequencies of the reference atlas as input for similarity assessment across the 13 cancer types (**Fig. 2A**). Samples of different tumor types intermixed and cancer type was not a main source of variance as shown in a t-distributed stochastic neighbor embedding (tSNE) of immune cell type proportions, (**Fig. 2B**). In contrast, abundance of Th17 T-cells explained most variance in the dataset (principal component 1, 43.7%), followed by the relative frequencies of M2 macrophages and terminally exhausted CD8 T cells (32.6% and 9.5%, respectively; **Fig. 2C**).

**Figure 2.**
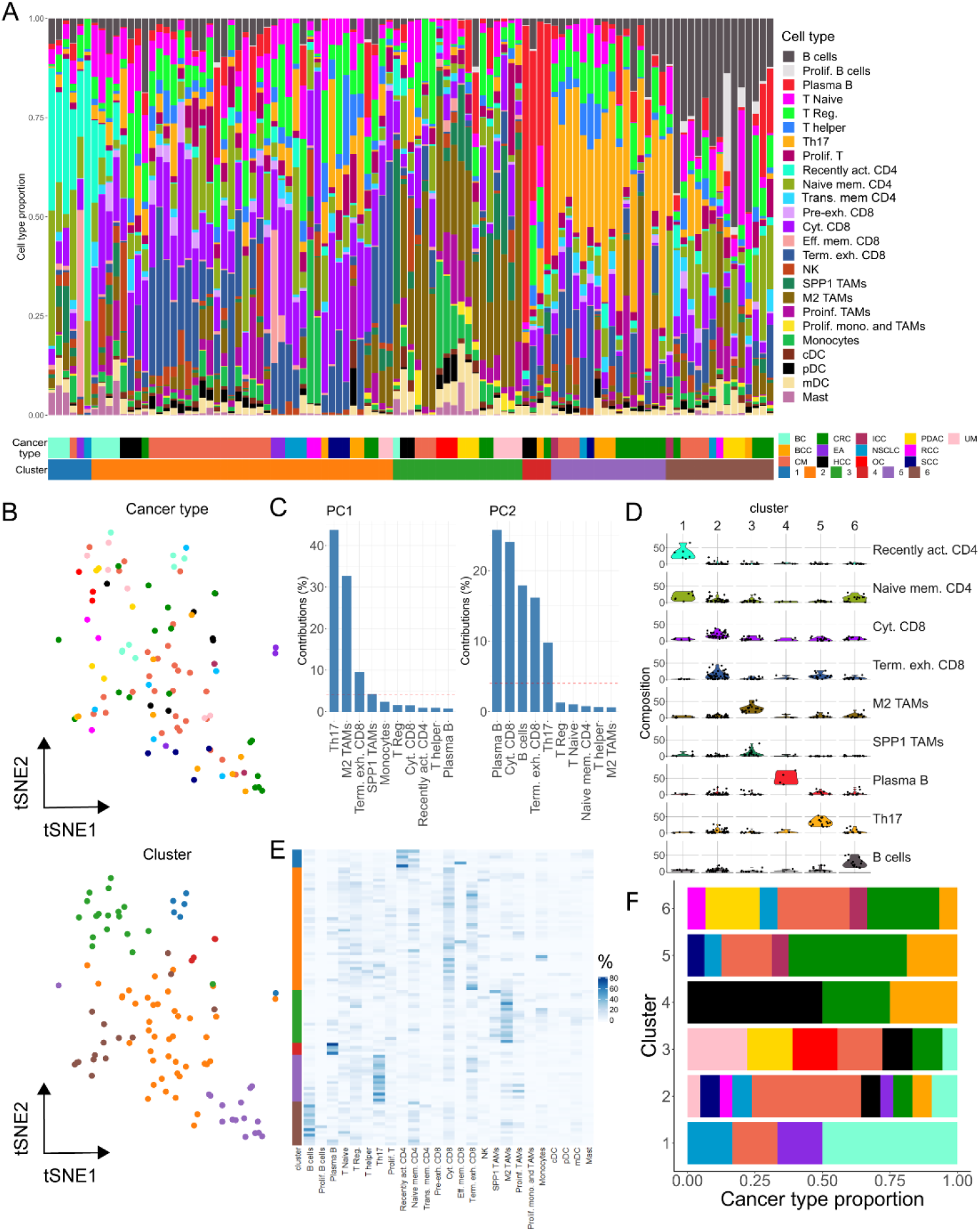
Patient stratification based on the tumor immune cell composition. **(A)** Cell type composition of patients colored by cell type/state frequencies. Patients are clustered (C1-6) into group of similar cell type composition. Cancer and cluster identities are indicated below. **(B)** Dimensionality reduction representation (tSNE distribution) of cell type frequencies in cancer patients colored by cancer type (top) and cluster identity (bottom). **(C)** Variance contribution to the two first principal components (PC) of the top variable cell types. **(D)** Frequencies (% total cells) of cell types representative for cluster 1-6. **(E)** Heatmap representation of cell type frequencies within each cluster. **(F)** Cancer type contribution to the six immune clusters; color code as cancer types in (A).

Correlating immune cell population frequencies identified an intriguing pattern of tumors being mutual exclusive for the presence of lymphoid or myeloid cell types (**Supplemental Fig. 5A**). Moreover, lymphoid-enriched tumors presented either naïve-memory, effector-memory and recently activated profiles or more differentiated (regulatory, cytotoxic and exhausted) phenotypes. Myeloid-enriched tumors further split in pro-inflammatory and inhibitory (M2 and SPP1 TAMs) subtypes, the former correlating with the presence of plasmacytoid and myeloid DCs. A hierarchical *k*-means clustering using immune cell proportions as features defined six clusters with largely different compositions (**Fig. 2A,B,D,E**). In spite of the different sizes of the clusters, almost all cancer types were presented in each cluster, confirming large commonalities of immune cell compositions between them (**Fig. 2F**). Tumors in cluster 1 (C1) showed a high proportion of recently activated and naive-memory CD4 T-cells, whereas C2 had high amounts of cytotoxic and terminally exhausted CD8 cells (**Fig. 2D,E**). C3 displayed exceptional high levels of macrophages (M2 and SPP1), C4 was driven by increased frequencies of plasma B-cells, C5 contained high proportions of Th17 T-cells, and C6 was very high on B-cells. Consistent with the distinct cell type proportions, the clusters presented specific gene expression signatures (**Supplemental Fig. 5B** and **Supplemental Table 4)**. Intriguing from a therapeutic perspective, C5 showed a striking increase of exhaustion markers on CD8 cells with significantly higher levels of *LAG3, PDCD1* and *CTLA4* (ANOVA, p<0.01; **Supplemental Fig. 5C,D**). The fact that these markers are also target for immune-therapy suggests C5 tumors to be more susceptible to ICI treatment, but could also shed light on the role of Th17 cells, strongly enriched in these patients (**Fig. 2D,E**).

To facilitate the classification of immune profiles of future datasets, we trained a RF classifier with the 25 immune cell population achieving a highly accurate classification (accuracy: 0.8, 95% CI: (0.593, 0.932), p-value: 5.36e-05, **Online Methods**). Of note, the cluster-specific cell types were also the most important variables when training the RF classifier using cell type proportions as features to predict cluster identities (**Supplemental Fig. 6**). We believe pan-cancer immune classifications to be highly informative and clinically predictive when integrating tumor samples from single-cell studies with patient survival and response to therapy meta-data. We envision that this classifier can be a useful future patient stratification tool for prognosis and immuno-therapy response. Moreover, using the classifier, the pan-cancer immune classification system could be extended to additional cancer types and drive the design of basket clinical trials in which a common immune stratification and recruitment framework is applied across cancer types.

### A resource for immune cell annotation

To demonstrate the predictive value of the atlas, we generated query datasets from different cancer types and varying experimental designs. Following clustering of the query datasets, we projected either single cells or clusters onto the atlas using a reference-based projection (Butler et al. 2018) or cluster matching (Mereu et al. 2020) tools, respectively. As proof-of-concept, we performed scRNA-seq for human primary OC and uveal melanomas (UM), liver metastases (from primary UM) as well as a brain metastasis (from primary CM).

Using cell-by-cell projection, we matched UM and OC cells to specific cell types and states of the atlas reference (**Fig. 3A,D**). Query cells projected to macrophage and T-cells, with a large variety of cell states being detected. The UM harbored all macrophage subtypes, including abundant SPP1+ and M2 TAMs and a small fraction of proliferative macrophages (**Fig. 3A**). In addition, the entire spectrum of T-cell phenotypes could be assigned, with a high fraction of cells projecting to the terminally exhausted CD8 T-cell state. The OC dataset contained less SPP1+ TAMs, and exhibited an increased proportion of proliferative macrophages (**Fig. 3D**). We also detected a small number of conventional DC and monocytes. We next used clustering analyses on the query datasets and subsequently matched the identified clusters to the atlas reference. There were 7 and 3 clusters for UM and OC, respectively (**Fig. 3B,E**). Matching the query and reference atlas clusters resulted in a clear separation between the lymphoid and myeloid populations (**Fig. 3C,F**). In UM, cytotoxic and terminally exhausted cells were enriched in cluster 1 and cluster 6, in line with their distribution in the dimensionally reduction representation (**Fig. 3A**). SPP1 TAMs showed higher scores in cluster 2, while M2 TAMs fell into cluster 4, an assignment consistent with the cell-by-cell projection. Cluster 7 matched to both lineages likely depicting cell doublets.

**Figure 3.**
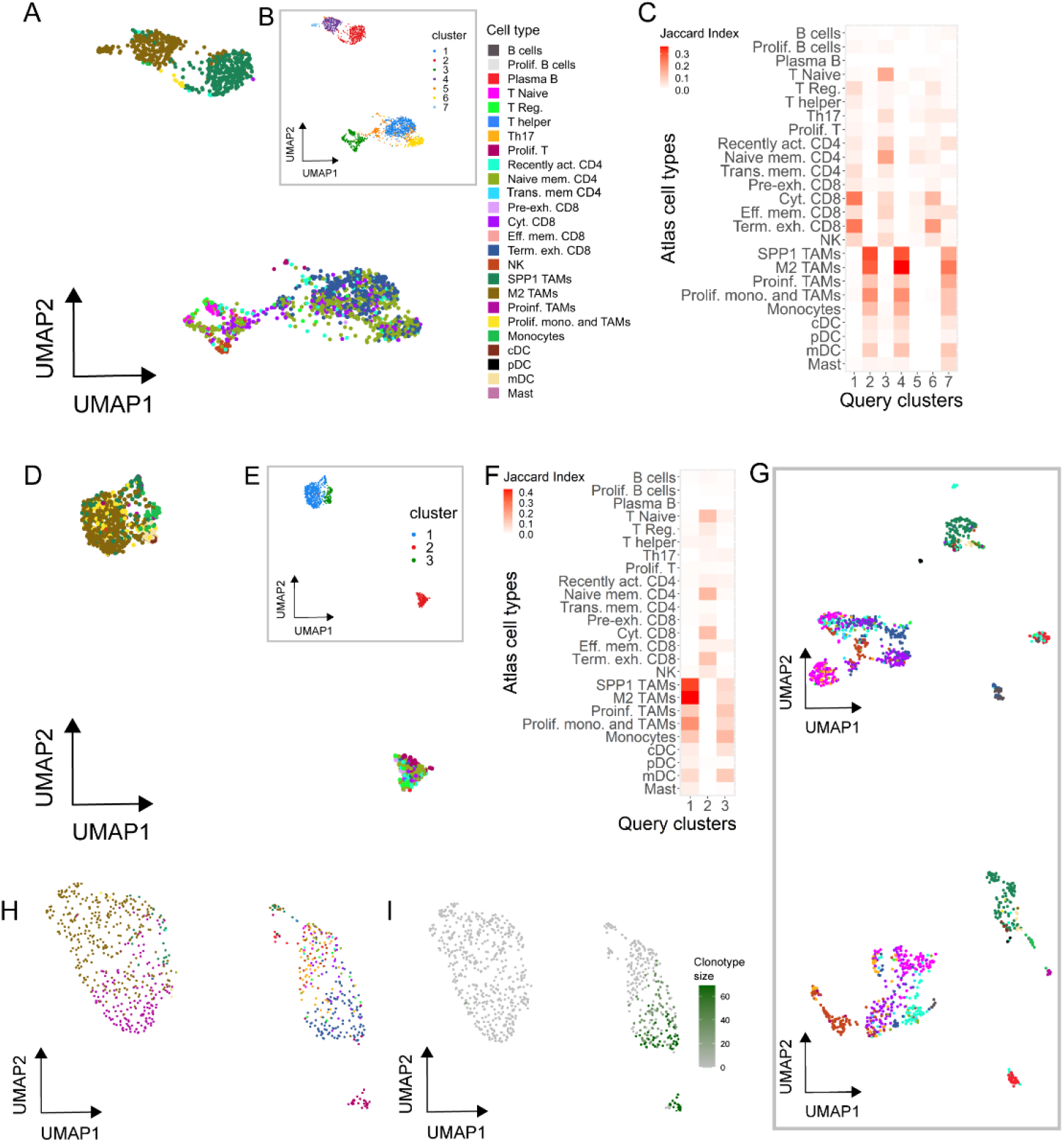
Automated annotation of external human tumor-derived immune cells using the tumor immune atlas as reference. **(A,B)** UMAP representation of immune cell transcriptomes from a primary uveal melanoma colored by their predicted cell type/state based on the tumor immune reference (**A**) or using unsupervised clustering (**B**). **(C)** Marker correspondence (Jaccard index) between uveal melanoma clusters (B) and the cell types clusters of the reference atlas. **(D,E)** UMAP representation of immune cell transcriptomes from a primary ovarian carcinoma colored by predicted cell type/state **(D,** color code as in A) and after clustering (**E**). **(F)** Marker correspondence (Jaccard index) between the ovarian cancer clusters and the cell type clusters of the reference atlas. **(G)** UMAP representation immune cells from two uveal melanoma liver metastasis colored by their predicted cell type (color code as in A). **(H,I)** UMAP representation of T-cells isolated from a brain metastasis colored by their predicted cell type (**H**, color code as in A) and clonal expansion profiled through TCR genotyping (**I**).

The projection of cells from UM liver metastases rendered a precise separation of T-cell states (**Fig. 3G**), which split into naïve, recently activated, cytotoxic and proliferative subpopulations. We found a distinct frequency of exhausted cells between both UM metastases. Although much less abundant, also all DC and B-cell states could be assigned in both samples. Cell-by-cell projection of T-cells from a NSCLC brain metastasis also identified most cell states, with a high proportion of terminally exhausted CD8 T-cells (**Fig. 3H**). Combined scRNA-seq and TCR genotyping assigned clonally expanded T-cells as being either actively proliferating or terminally exhausted, in line with the expected history of tumor reactivity and proliferation of exhausted cells (**Fig. 3H,I**) (Azizi et al. 2018; Li et al. 2019).

We further wondered about the applicability of the atlas as reference across species. To tackle this question, we generated scRNA-seq datasets for two liver metastases derived from mouse CRC organoids (Tauriello et al. 2018), one CD45 selected and the other enriched in T-cells, and projected individual cells onto the human reference. Intriguingly, the mouse datasets could be projected with high confidence (~70% of cells with matching probabilities >0.5). Mouse clusters of main subtypes could be readily annotated and specific subpopulations could also be assigned to distinct T/B-cell or macrophage cell states using the human reference (**Figure 4A-D**). Again, distinct T-cells states emerged from the dimensional reduction plots or after clustering analysis (**Figure 4A-C**). Consistently, cell projection of the experimentally enriched T-cell fraction to the reference atlas identified defined T-cell states (**Figure 4E**). Clonal expansion according to TCR sequencing could only be detected in cells assigned to be proliferative, cytotoxic or exhausted, while naïve, naïve-memory and recently activated cells were not of clonal origin (**Figure 4E,F**).

**Figure 4.**
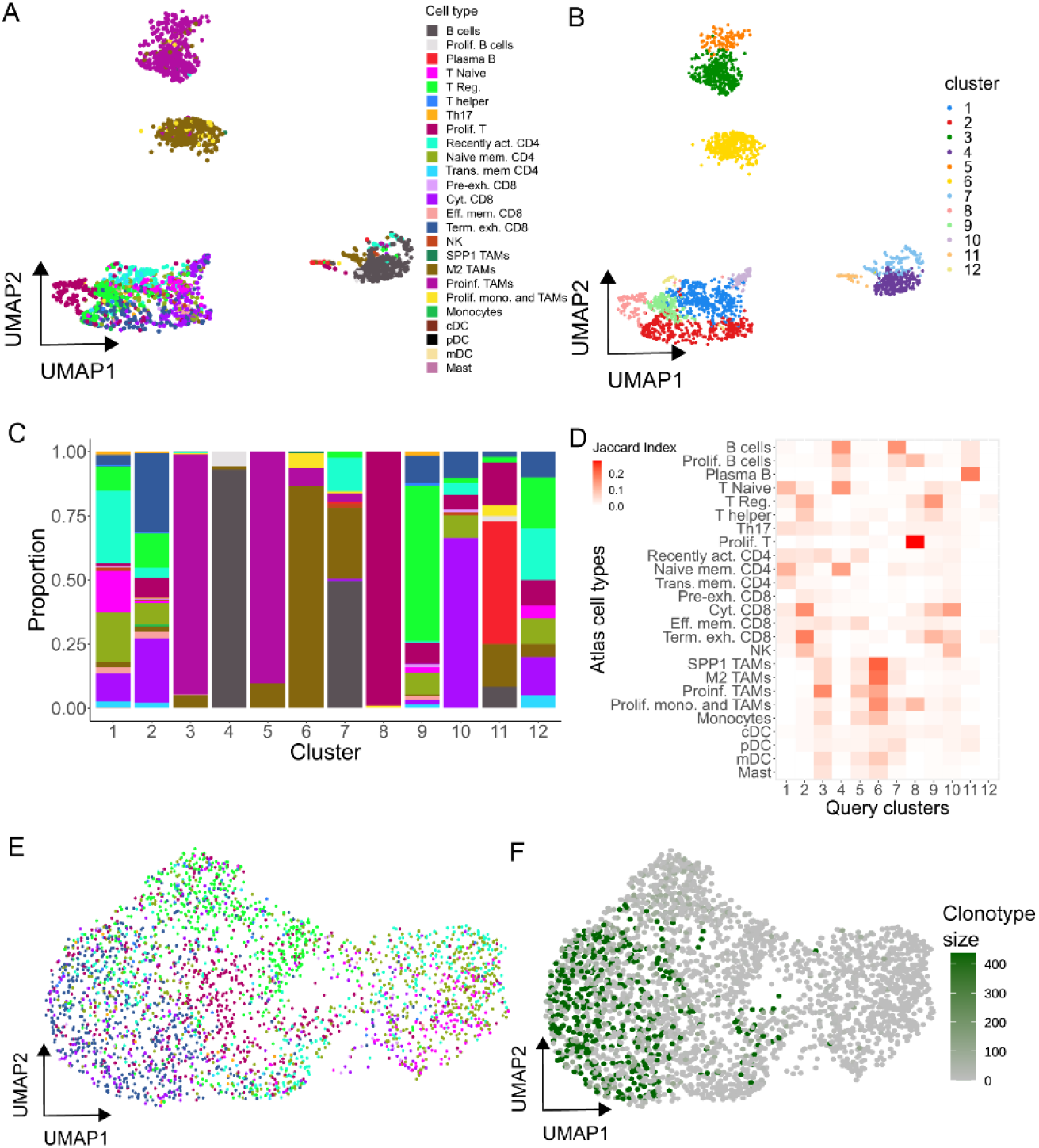
Projecting mouse tumor immune cells onto the human reference atlas. **(A,B)** UMAP single-cell transcriptome representations of mouse T-cells isolated from tumor organoids. Cells are color-coded by their predicted cell type based on the human reference atlas (**A**) or by cluster identity (**B**). **(C)** Cell type composition of each cluster (color code as in A). **(D)** Marker gene correspondence (Jaccard index) between the mouse immune clusters and the cell type clusters of the reference atlas. **(E,F)** UMAP representation of mouse T-cells isolated from tumor organoids colored by predicted cell type (**E**, color code as in A) and clonality based on expanded TCR clonotypes (**F**).

### Spatial localization of immune cells in tumor sections

Widespread evidence indicates that the spatial distribution of immune cells is important for ICI response (Chen and Mellman 2017). To explore this issue, we combine single-cell reference atlas immune profiles with ST data from tumor sections. The aim of this approach is to provide spatial tumor maps that precisely delineate co-localization of immune cells in tumors. To integrate both data modalities, we applied *SPOTlight*, a non-negative matrix factorization (NMF) based spatial deconvolution framework (Elosua et al. 2020). *SPOTlight* identifies cell type-specific topic profiles (gene expression signatures) from scRNA-seq data, which are subsequently used to deconvolute ST spots (**Supplemental Fig. 7**). To predict and quantify the location of the 25 cell types and states of the tumor immune reference, gene expression profiles were translated into *SPOTlight* topic profiles. Topics were highly specific to clusters resulting in high specificities to localize immune cell types (**Figure 5A**). To map immune cells within tumor sections, we generated ST datasets for a oropharyngeal squamous cell carcinoma (SCC) metastases and analyzed two sequential sections of an invasive ductal BC.

**Figure 5.**
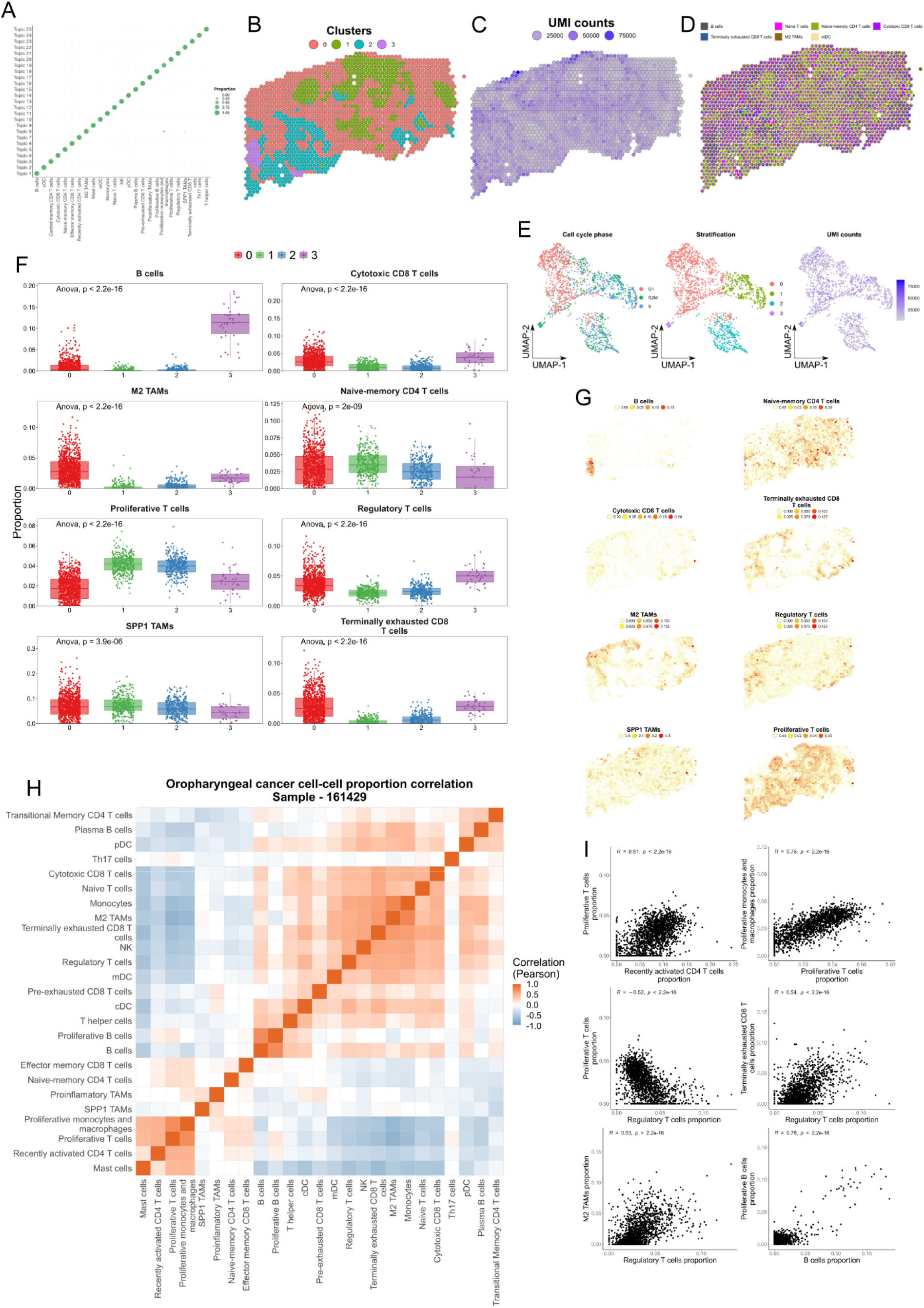
Spatial mapping of the reference immune cell types using ST oropharyngeal squamous cell carcinoma (SCC) sections. **(A)** Cell type specific topic profiles presenting a high topic/cell type specificity. **(B)** ST profiled section of a SCC primary tumor. Tissue stratification according to unsupervised clustering. **(C)** The number of unique molecular identifiers (UMI) recovered from each spot indicate the areas transcriptionally most active. **(D)** Pie chart representation showing proportions (per ST spot) of *SPOTlight* predicted immune cells based on the single-cell immune reference atlas. To visualize spatially variable cell types, only immune cell types present in <90% of the spots are displayed. **(E)** UMAP embedding of ST spots presenting the cell cycle phase (left), cluster identity (middle), and UMI counts (right) for each spot. **(F)** Box plots displaying significant differences in cell type proportion of clusters (ANOVA test). **(G)** Location and proportion of significantly differentially located cell types in the SCC section. **(H)** Clustered correlation matrix between the predicted cell type proportions identifying co-localization (red) and exclusive (blue) immune distribution patterns. **(I)** Scatter plots of (anti-) correlating cell type pairs identified in (H).

Clustering the spatial SCC data identified four distinct transcriptional and proliferative active areas containing cancer cells (cluster 1/2), surrounded by stroma (cluster 0) and an area enriched in immune cells (cluster 3, **Figure 5B-E**). Mapping the immune cell distribution in the sections using *SPOTlight,* returned proportions for each cell within a spot. This resulted in a clear regionalization of distinct immune cell types and states (**Figure 5D**, **Supplemental Fig. 8**). The cancer areas cluster 1/2 presented a similar immune infiltration pattern, with an enrichment of proliferative T-cells and SPP1 macrophages (**Fig. 5F,G**, **Supplemental Fig. 9**). Naive-memory CD4 T-cells were further enriched in the cluster 1 region. Cluster 3, in turn, presented a distinct immune infiltration pattern characterized by an enriched presence of (proliferative) B-cells. The stroma section, cluster 0, harbored regulatory T-cells and terminally exhausted CD8 T-cells and was specifically enriched in M2 macrophages and naive T-cells. An interaction matrix computed based on the predicted proportions on the spatial distribution of cell states in the section highlighted co-localization and mutual exclusive patterns (**Figure 5H,I**). Here, spatial co-localization provided further insights into immune-modulating mechanisms of cancer and immune-suppressive stromal cells (e.g. regulatory T-cells). Exemplarily, proliferative T-cells co-localized with recently activated, effector/naive memory T-cells and Th17 T-cells, while they were excluded from areas with regulatory, cytotoxic and exhausted T-cells as well as M2 TAMs. Accordingly, regulatory T-cells showed a positive correlation with cytotoxic cells (including NK), pre- and terminally exhausted CD8 T-cells and M2 TAMs, but a negative correlation with proliferating and activated T-cells. It has been shown that regulatory T-cells inhibit the proliferation of effector T-cells by releasing CD39 and CD73 that bind to the A2A receptor (Allard et al. 2020). In line with this finding, and in addition to the exclusive pattern of regulatory and proliferating T-cells, we detected a positive expression correlation of CD39 *(ENTPD1)* and CD73 *(NT5E)* with the regulatory T-cell predicted proportions (p <2.2e^-16^ and p=2.5e^-06^ respectively) and an anticorrelation with proliferating T-cells (p <2.2e^-16^ and p=9e^-09^ respectively). Along with the detected spatial organization with other immune cell types, regulatory T-cells illustrate the value of spatial immune mapping to advance our understanding of tumor immunology. Analysis of a second SCC case confirmed the regional restriction of tumor immune cells and confirmed the previously described co-localization and mutually exclusive patterns (**Supplemental Figs. 10-12**).

A striking cancer-specific regional distribution was also observed in the invasive ductal BC and replicated over serial sections of the tumor (**Figure 6A-C** and **Supplemental Fig. 13**). The gene expression profiles of cancer cells pointed to a strong sub-clonal structure, with mutually exclusive localizations of BC subtype-specific HER2+ (ERBB2) and ER+ (ESR1) clones. Intriguingly, the subclonal architecture was directly associated with local enrichment of distinct immune cell states (**Figure 6D** and **Supplemental Fig. 14,15**). The ESR1+/HER2-region showed increased presence of proliferative T-cells (**Figure 6E,F** and **Supplemental Fig. 14,15**). In contrast, the HER2+/ESR1-region showed a higher proportion of regulatory and cytotoxic T-cells, M2 TAMs and NK cells. In this region, coupled to the higher proportion of regulatory T-cells, we also observed an enrichment of pre-exhausted CD8 T-cells, suggesting incipient inactivation of the immune cell response. Lastly, we observed an enrichment of plasma B-cells and SPP1 TAMs lining the cancer areas in the fibrotic tissue. The importance of a quantitative assessment of cell types within ST spots is displayed by the ubiquitous presence of proliferative and helper T-cells across all areas, but the significant proportional enrichment in the tumor fraction (**Figure 6F**). Intriguingly similar to the SCC, spatial correlation analysis in BC, among other features, identified the co-localization of proliferative T-cells and macrophages and the anti-correlation with regulatory and cytotoxic T-cells and M2 TAMs (**Figure 6G,H**). The tumor immune cell reference, in combination with *SPOTlight*-based spatial mapping of ST data, enables to harmonize the spatial assessment of tumor infiltrating immune cells; a first step towards an automated digital pathology framework. We foresee the regional distribution of immune cell types to become an important feature for the prediction of immuno-therapy outcome. Moreover, longitudinal sampling for spatial immune cell mapping throughout ICI treatment could aid explaining therapy action and point to further weaknesses approachable through combinatorial therapy strategies.

**Figure 6.**
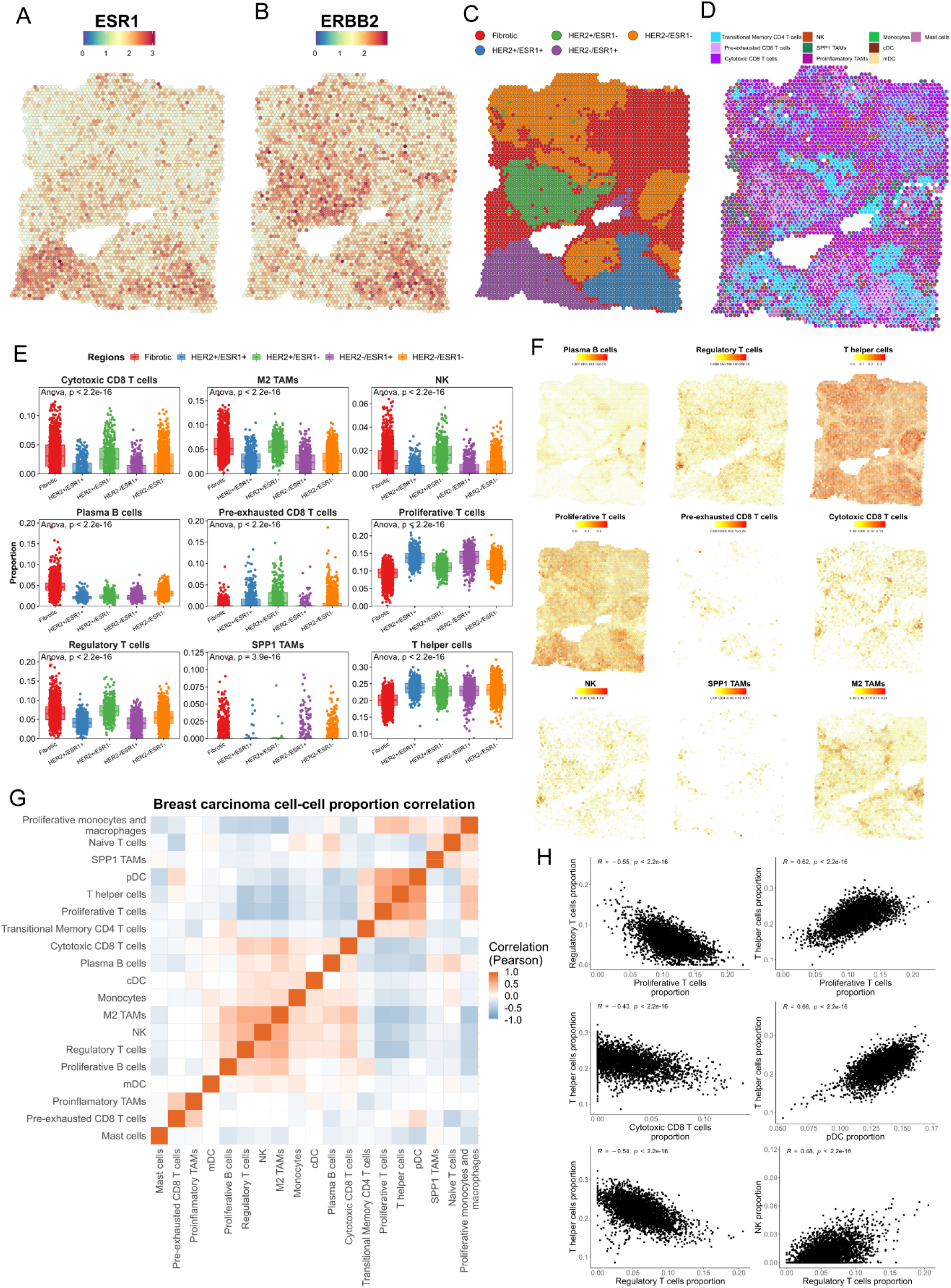
Tumor immune reference mapping ST section from a ductal breast carcinoma. **(A,B)** Estrogen receptor (ESR1, **A**) and HER2 receptor (ERBB2, **B**) gene expression levels on the ST section, indicating profound regionality of the expression. **(C)** Tissue stratification and labeling according to unsupervised clustering. **(D)** Pie chart representation showing proportions (per ST spot) of *SPOTlight* predicted immune cells based on the single-cell immune reference atlas. To visualize spatially variable cell types, only immune cell types present in <90% of the spots are displayed. **(E)** Box plots of significantly differentially localized cell type proportions between the clusters (ANOVA test). Differences between tumor areas (i.e. HER+ and ER+) are observed, suggesting differential tumor microenvironments of the tumor subclones. **(F)** Location and proportion of immune cell types with local enrichment **(G)** Clustered correlation matrix between the predicted cell-type proportions providing evidence for immune cell type co-localization in tumors. **(H)** Scatter plots of proportions of correlating cell type pairs (G).

## Conclusion

Our atlas of 317,111 immune cells from 13 cancer types allowed us to catalogue tumor-resident immune cells across a large patient cohort and to draw a comprehensive tumor immune cell atlas of human cancers. We envision the application of the atlas in multiple areas of precision oncology. Therefore, we provide a framework for an immune-based patient stratification, the feasibility to integrate newly generated patient single-cell data and a toolbox to map immune cells directly in tumor sections. Following the integration of clinical cohort single-cell studies with patient outcome and response meta-data, we expect the atlas to be predictive for patient prognosis and immuno-therapy response to a level that greatly exceeds currently applied stratification strategies.

## Acknowledgements

This publication is part of a project (BCLLATLAS) that has received funding from the European Research Council (ERC) under the European Union’s Horizon 2020 research and innovation programme (grant agreement No 810287). This work has received funding from the Ministerio de Ciencia, Innovación y Universidades (SAF2017-89109-P; AEI/FEDER, UE) and the Fundació La Marató de TV3 (201903-3031-32). We further acknowledge funding from the St Vincent’s Clinic Foundation (VC) and the National Health and Medical Research Council Investigator Grant (APP1175781, JEP), the Fundación Asociación Española contra el Cáncer (AECC), FERO (EDM), Ramón Areces Foundation, Cellex Foundation, BBVA (CAIMI), the ISCIII, FIS (PI16/01278), Juan de la Cierva formación fellowship (CRP) and Sara Borrell fellowship (EPR). Core funding is from the ISCIII and the Generalitat de Catalunya. We acknowledge support of the Spanish Ministry of Economy, Industry and Competitiveness (MEIC) to the EMBL partnership, the Centro de Excelencia Severo Ochoa, the CERCA Programme / Generalitat de Catalunya, the Spanish Ministry of Economy, Industry and Competitiveness (MEIC) through the Instituto de Salud Carlos III and the Generalitat de Catalunya through Departament de Salut and Departament d’Empresa i Coneixement. We also acknowledge the Co-financing by the Spanish Ministry of Economy, Industry and Competitiveness (MEIC) with funds from the European Regional Development Fund (ERDF) corresponding to the 2014-2020 Smart Growth Operating Program.

## Author Contributions

HH designed the study. PN and MEB performed all data analyses. JL analyzed TCR data. EM and CRP supported the data analysis. DM, CM, SR, PL and EP generated single-cell datasets. DK, CC generated ST datasets. MS and AH supported data interpretation. VTC, RG, IG, JMP, JS, JP and EB provided samples, technical and sequencing support. HH, PN and MEB wrote the manuscript with contributions from the coauthors. All authors read and approved the final manuscript.

## Data availability

The Tumor Immune Cell Atlas count matrix is deposited at https://zenodo.org/record/4036020#.X2ijWWgzZEY. The analysis code is available under https://github.com/Single-Cell-Genomics-Group-CNAG-CRG/Tumor-Immune-Cell-Atlas. The atlas raw data is available as stated in the Online Methods. To explore the atlas, to project external data and to apply the immune classifier, we are currently developing a user-friendly ShinyApp (in progress). Raw data for the projected single-cell datasets and ST will be available at the Gene Expression Omnibus (GEO) database under GSE158803.

## Online Methods

### Tumor sample preparation

Human brain metastasis of cutaneous melanoma (CM) patients were diagnosed and treated at the Vall d’Hebron University Hospital (Barcelona, Spain). A written consent, indicating that the samples obtained as standard of care in the clinical practice could be used for research, was obtained from the parents/legal representant in agreement with the declaration of Helsinki. The study was approved by the local IRB. The uveal melanomas (UM), liver metastasis of the uveal melanoma and ovarian cancer (OC) were obtained from the Catalan Institute of Oncology (ICO). The study was approved by the local IRB: PR167/18 – Identification of immunity against GNAQ/11 mutations in uveal melanoma patients (PR070/18 – Identification of the antigenic potential of driver mutations in Metastatic Uveal Melanoma. Generating therapeutic windows of opportunity).

After tumor resection, tissues were enzymatically digested using human tumor dissociation kit (Miltenyi Biotec). To isolate CD45+ cells, CD45 TIL microbeads (Miltenyi Biotec) were used following manufacturer’s instructions and stored in cold 1X PBS supplemented with 0.1% BSA until partitioning for single cell RNA-sequencing. The uveal melanoma and ovarian cancer samples were transferred from the surgery room to the single-cell laboratory in DMEM medium (Gibco) on ice (<1h). Upon arrival to the laboratory, samples were washed twice with cold 1X HBSS (ThermoFisher Scientific) and minced on ice using razor blades. Next, the samples were incubated during 15 min at 37°C under constant rotation at 700 rpm with 2 ml of a pre-warmed dissociation mix (200 U/ml Collagenase IV, 1X HBSS). Every 5 min, samples were pipetted with a wide bore p1000 tip to help tissue dissociation. After 15 min, samples were passed 10 times through a 0.9 mm needle and 10 additional times through a 0.6 mm needle. Then, the enzymatic digestion was stopped by adding 10% fetal bovine serum (FBS). Samples were filtered with a 70 μM cell strainer (pluriSelect) and centrifuged during 5 min at 500 rcf at 4°C. Pellets were washed twice with cold 1X HBSS and resuspended in 1X PBS supplemented with 0.05% BSA (Miltenyi Biotec). Cell concentration and viability were verified by counting with a TC20™ Automated Cell Counter (Bio-Rad Laboratories, S.A). In the case of UM liver metastasis, CD45+ cells were isolated by Magnetic-activated cell sorting (MACS) using the OctoMACS™ Separator and MS columns (Miltenyi Biotec) following manufacturer’s instructions. Briefly, cells were incubated with human CD45 MicroBeads during 30 min at 4°C and CD45+ cells were separated by applying the cell suspension onto a pre-rinsed MS column. Cell concentration and viability were again verified with a TC20™ Automated Cell Counter (Bio-Rad Laboratories, S.A).

For the ovarian cancer, cells were subjected to a Cell Hashing protocol before proceeding to single-cell RNA sequencing. Cell hashing was performed following manufacturer’s instructions (Cell hashing and Single Cell Proteogenomics Protocol Using TotalSeq™ Antibodies; BioLegend). Briefly, the sample was resuspended in Cell Staining Buffer (Bio Legend) and split into four tubes with an equal number of cells. Next, sample tubes were incubated during 10 min at 4°C with Human TruStain FcX™ Fc Blocking reagent (BioLegend) and then a specific TotalSeq-A antibody-oligo conjugate was added to each tube and incubated on ice for 1h. Cells were then washed three times with cold PBS-0.05% BSA (ThermoFisher) and centrifuged for 5 min at 500 rcf at 4°C. Finally, cells were resuspended in an appropriate volume of 1X PBS-0.05% BSA in order to obtain a final cell density > 500 cells/ul, suitable for 10x Genomics scRNA-seq. An equal volume of hashed cell suspension from each of the four tubes was mixed and filtered with a 40 μm strainer. Cell concentration was verified by counting with a TC20™ Automated Cell Counter (BioRad Laboratories, S.A).

### Mouse Colorectal cancer (CRC)

Isolation of tumor cells for generation of mouse tumor organoids (MTOs) and in vitro expansion was previously described (Tauriello et al. 2018). Briefly, for mouse liver metastasis generation, 3-day grown MTOs were trypsinized followed by mechanical dissociation to obtain a single cell suspension. 6-8-week-old C57BL/6J mice (Janvier Labs) were subjected to intrasplenic injection of 3×10^5 single cells in 70μl HBSS, followed by splenectomy to prevent splenic tumor growth. The mice were euthanized 16 days postinjection and livers were collected in HBSS.

Tumor nodules were dissected from livers and minced with scalpels. The tissue was enzymatically digested with 0.2 mg/ml Collagenase IV (Sigma, ref: C5138), 0.2 mg/ml Dispase II (Sigma, ref: D4693) and 0.04 mg/ml DNase I (Sigma, ref: 10104159001) in 10%FBS/DMEM (Life Technologies). Enzymatic digestion consisted of mechanical dissociation during 4 cycles of 20 minutes at 37°C. The enzymatic reaction was quenched at the end of each cycle, by the addition of 30 ml of ice-cold HBSS (10% FBS supplemented). The cell suspension was filtered using 100 μm strainers (Corning). Lysis of erythrocytes was performed using red cell lysis buffer (150 mM NH4Cl, 10mM NaHCO3, 0.1 mM EDTA). Cells were incubated with FACS buffer (PBS, 2% FBS) containing blocking antibodies against CD16/CD32 at 4°C for 20 minutes to block the Fc receptor. Then, BV605-conjugated antibodies against EpCAM were used to stain the cells for 20 minutes at 4°C. Lastly, EpCAM-negative cells were sorted in a FACS Aria Fusion flow cytometer (BD Biosciences) from total viable cells, determined by nuclear staining with DAPI (Sigma).

### Single-cell RNA sequencing (scRNA-seq)

Cells were partitioned into Gel Bead-In-Emulsions by using the Chromium Controller system (10x Genomics) aiming at Target Cell Recovery of 5000-7000 total cells per sample, in the case of CM, UM, and mouse CRC, and a Target Cell Recovery of 10,000 total cells in the case of the OC. For the CM brain metastasis and a mouse CRC sample, single-cell Gene Expression (GEX) and T cell receptor (TCR)- enriched libraries were prepared using the Chromium Single Cell 5’ Library and Gel Bead Kit (10x Genomics, Cat. N. 1000006) following manufacturer’s instructions. In the case of the UM, OC samples and one mouse CRC, gene expression libraries were prepared using the Chromium Single-cell 3’ mRNA kit (V3; 10X Genomics, Cat. N. 1000075), following manufacturer’s instructions. Briefly, after GEM-RT clean up, cDNAs were amplified during 16 cycles for CM and mouse CRC samples, 12 cycles for UM samples and 11 cycles for the ovarian cancer sample. cDNA QC and quantification were performed on an Agilent Bioanalyzer High Sensitivity chip (Agilent Technologies).

For the OC sample, the GEX library was prepared using the single-cell 3’ mRNA kit (V3; 10x Genomics) with some adaptations for cell hashing, as indicated in TotalSeq™-A Antibodies and Cell Hashing with 10x Single Cell 3’ Reagent Kit v3 3.1 Protocol by BioLegend. Briefly, 1 μl of 0.2 μM HTO (Hashtag Oligonucleotides) primer (Integrated DNA Technologies, IDT) was added to the cDNA amplification reaction in order to amplify the hashtag oligos together with the full-length cDNAs. A SPRI selection cleanup was done in order to separate mRNA-derived cDNA (>300 bp) from antibody-oligo-derived cDNA (<180 bp), as described in the above-mentioned protocol from BioLegend. The cDNA sequencing library was prepared following 10x Genomics single-cell 3’ mRNA kit protocol, while HTO cDNA was indexed by PCR as follows. Briefly, 5 μl of purified hashtag oligo cDNA were mixed with 2.5 μl of 10 μM Illumina TruSeq D708_s primer (IDT), 2.5 μl of SI primer from 10 X single-cell 3’ mRNA kit, 50 μl of 2X KAPA HiFi PCR Master Mix (KAPA Biosystem) and 40 μl of nuclease-free water. The reaction was carried out using the following thermal cycling conditions: 98°C for 2 min (initial denaturation), 12 cycles of 98°C for 20 sec, 64°C for 30 sec, 72°C for 20 sec, and a final extension at 72°C for 5 min. The HTO library was purified by adding 1.2X SPRI select reagent to the PCR reaction.

All cDNA libraries were indexed by PCR using the PN-220103 Chromiumi7 Sample Index Plate. Size distribution and concentration of 3’ and 5’ GEX libraries, TCR-enriched and HTO libraries, were verified on an Agilent Bioanalyzer High Sensitivity chip (Agilent Technologies). Finally, sequencing of all libraries was carried out on an Illumina NovaSeq6000 sequencer to obtain approximately 40,000 reads/cell, in the case of GEX libraries, and 2000 reads/cell for the TCR-enriched and HTO libraries.

### Antibodies hashtag oligo sequences

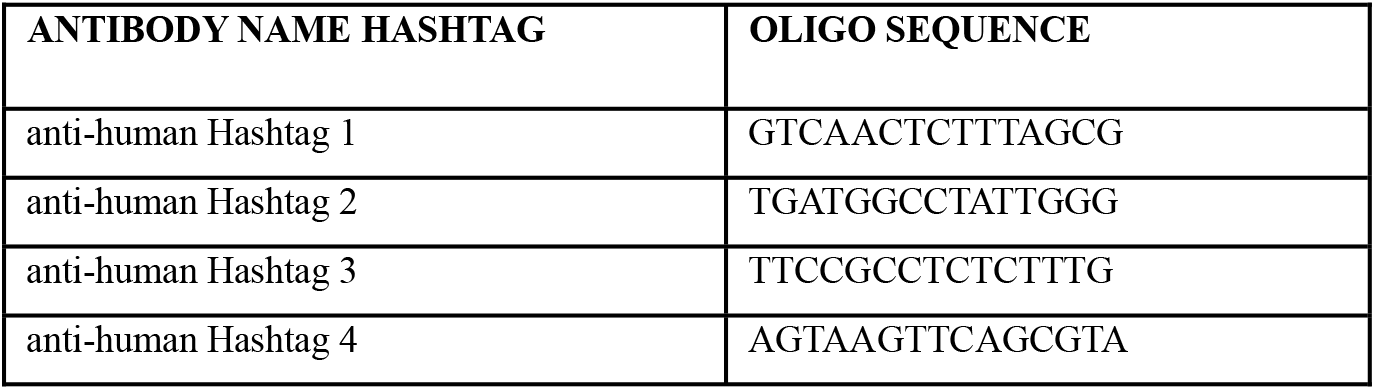

### Primers used for single cell library construction

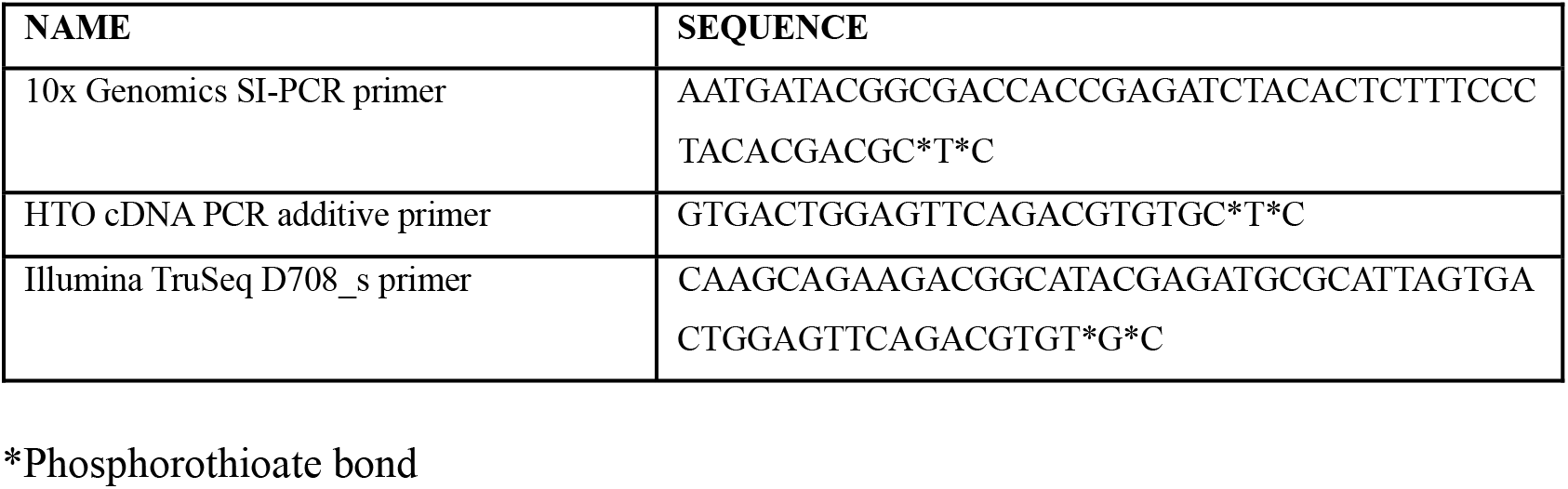

### Preprocessing the data

We initially pre-processed each scRNA-seq dataset independently: breast carcinomas (BC, GSE114727, GSE114725), basal cell (BCC) and squamous cell carcinomas (SCC, GSE123814), endometrial adeno- (EA) and renal cell carcinomas (RCC, GSE99254), intrahepatic cholangio- (ICC) and hepatocellular carcinomas (HCC, GSE125449, GSE140228), colorectal cancers (CRC, GSE132465, GSE99254), pancreatic ductal adenocarcinomas (PDAC, GSA CRA001160), ovarian cancers (OC, https://figshare.com/s/711d3fb2bd3288c8483a), non-small-cell lung cancers (NSCLC, E-MTAB-6149, E-MTAB-6653, GSE97168, GSE99254), and cutaneous (CM, GSE120575, GSE123139) and uveal (UM, GSE139829) melanomas.

From the original count matrices, we filtered the low-quality cells based on mitochondrial RNA percentage, number of UMIs (unique molecular identifiers) and number of different genes (thresholds adjusted separately for each dataset). Then, we obtained the highly variable genes, corrected, logarithmically normalized and scaled the counts (default parameters) of each dataset using Seurat (version 3.2.0) (Stuart et al. 2019). Principal Component Analysis (PCA) was performed on all individual datasets. The optimal number of principal components ranged between 25 and 40, depending on dataset size. Dimensionality reduction was performed by applying the Uniform Manifold Approximation and Projection (UMAP) algorithm, which also served as a two-dimensional embedding for data visualization. Cluster analysis was performed using the Louvain clustering algorithm, a method separating different communities inside the data by optimizing the modularity measure. The clusters were annotated by expression of canonical markers of major cell types to consistently identify and extract immune cells. Before dataset integration, the major cell types were identified, i.e. T-, B-, myeloid and dendritic cells and separated from the tumor cells. Note that for some datasets prior information for cell-level annotation was also available.

### Construction of the tumor immune cell atlas

To construct the atlas, we used the raw counts from the previously annotated immune cells of each dataset and followed Seurat’s standard data integration pipeline. This method is based on the identification of *anchor* cells between pairs of datasets, which are used to *harmonize* the integrated datasets. To do this, Canonical Correlation Analysis (CCA) is applied, for which we set the number of anchors to 3000. Prior to integration, we obtained the dataset-specific genes in order to remove them from the integration anchors, thereby reducing noise. To do this we joined all the datasets and with the *FindAllMarkers* function calculated the genes specific for each dataset. Then, we removed these genes from the integration features. Integrating multiple datasets is a computationally intensive procedure. Due to the large number of atlas cells, we used the alternative reciprocal PCA (RPCA) algorithm for anchor identification, as proposed by Stuart, Butler, et al. (Stuart et al. 2019). This method is notably faster and performs considerably similar to the standard CCA method. With this procedure, we built a tumor immune cell atlas of over 300,000 immune cells across 13 cancer types and 217 patients. After integration, we computed the clusters again to obtain a fine-grained resolution of cell types and states. We increased the resolution parameter in Seurat’s *FindClusters* up to 1.2, to retain more subtle differences between subpopulations.

We performed Differential Expression Analysis (DEA) for all clusters to determine their marker genes. For the DEA we used the normalized RNA counts, instead of using the integrated data, to increase the accuracy identifying significant cluster makers. To further refine the cluster signatures, we computed the clusters using both the Wilcoxon Rank Sum test and MAST (Finak et al. 2015) methods and kept only genes identified with both approaches. We annotated the refined clusters using a curated gene signature of immune cell subtypes and comparing it with the results of the DEA analysis (using the *matchSCore* package (Mereu et al. 2020)). Here, not only the major immune cell types, but also different immune cell states were identified.

### Validation of clusters and signatures using a random forest (RF) classifier

To assess the robustness of the clusters and their respective signatures, we trained a RF classifier as follows. First, we computed 25 cell type-specific signatures with the *AddModuleScore* function from *Seurat* using the markers in **Supplementary Table 3** that were kept in the integrated expression matrix. Additionally, we calculated a random signature per cell type by sampling without replacement as many genes as present in the real signatures. Second, we performed 5-fold cross validation to test both the bias and the variance of the classifier. We sought to ensure that all cell types were present in the five test sets. Thus, each test set of each fold was built by taking 20% of the cells in each cell type, which resulted in each cell appearing in one test set and four training sets. Third, we trained a RF classifier in each of the five training sets with the *randomForest* v4.6.14package; using the signatures as features and the cell type annotation as target variable. Finally, we tested the accuracy of the classifier in the respective test set using the *confusionMatrix* function from the *caret* v6.0.86 package.

### Clustering and classification of patients based on immune cell compositions

In order to find similarities in patients’ immune microenvironment across cancer types, we applied patient clustering using the proportion of 25 immune cell states of their TME. For this we filtered out patients with less than 500 cells and applied hierarchical K-means (HKM) clustering (Chen et al. 2005; Alashwal et al. 2019) using the cell types as variables and their proportion in the TME as values. We set the number of clusters to k = 6 (**Supplementary Fig. 5b**) and used the agglomerative Ward 2 method with Euclidean distance metric. This algorithm iteratively joins instances in clusters, reducing intra-cluster variance. Further, we built a RF classifier to predict the assignment of each patient to one of the six immune subtype clusters. Therefore, we split the dataset into training and test sets (75% and 25% of the patients, respectively) and grew 100 trees to train the classifier, while automatically assessing predictor’s importance and calculating the proximity between patients for a better model. To evaluate the variable importance obtained from the RF classifier, we assessed how the accuracy of the model decreased when a variable is excluded from the model (Mean Decrease Accuracy) and how each variable affects the homogeneity of the trees in the forest (Mean Decrease Gini).

### Dataset projection onto the tumor immune cell atlas

For the projection and annotation of external datasets onto the atlas, we used Seurat’s anchor-transferring method. This algorithm uses the PCA structure of the reference atlas and projects it on the query. Then it finds pairs of anchor genes between datasets that enable the projection. These anchors allow the transfer of annotations (labels) between datasets. Here, we used an unsupervised anchoring based on the first 30 principal components of the datasets and the RNA assays with a *LogNormalize* normalization method. We obtained the cell-type assignment of each cell in the query datasets plus the corresponding prediction probability. We also used *matchSCore2’*s function to correlate the markers of query clusters with the markers of the 25 cell types/states of the atlas. Therefore, we used the top 100 markers (average log-fold change) for each of the clusters.

### Human oropharyngeal cancer samples

All patients provided informed consent for the collection of human specimens and data. This was approved by the St Vincent’s Hospital Research Office (2019/PID04335) in accordance with the National Health and Medical Research Council’s National Statement of Ethical Conduct in Human Research. Patients undergoing surgical resection for a locally advanced oropharyngeal cancer were recruited to the study. After surgical removal, the anatomical pathologist dissected a sample of both the primary and nodal metastasis. Samples were tumour banked in accordance with our ethically approved protocol. Within 30 min of collection, tumour samples were tumour banked. Samples were cut into 1mm x 1mm chunks with a scalpel blade. For Visium, a tissue chunk was snap frozen in OCT. After freezing, samples were moved to liquid nitrogen for long term storage.

### Visium Spatial Gene Expression

Frozen tissue samples were processed using the Visium Spatial Gene Expression slide and reagent kit (10x Genomics, US) following the manufacturer’s instruction. Briefly, 10 μm sections were placed into the capture areas of the Visium slide. Tissue morphology was assessed with H&E staining and imaging using a Leica DM6000 microscope equipped with a 20x lens (Leica, DE). The imaged sections were then permeabilized for 12 minutes using the supplied reagents. The permeabilization condition was previously optimised using the Visium Spatial Tissue Optimisation slide and reagent kit (10x Genomics, US). After permeabilization, cDNA libraries were prepared, checked for quality and sequenced on a NovaSeq6000 platform (Illumina, US). Around 300 million pair-ended reads were obtained for each tissue section. Read 1, i7 index and Read 2 were sequenced with 28, 8 and 98 cycles respectively.

### Visium data quality control

Quality control was carried looking at the number of UMIs, genes and mitochondrial percentage. Spots with <1.000 UMIs were removed from the analysis due to poor quality. Three tissue slices were found on the same Visium slide area, 2 were mostly on the spots while the 3^rd^ was mainly off the capture area. The latter was discarded while the other 2 where separated and treated as different datasets. Data was scaled and normalized using *SCTransform* with default parameters. Principal component analysis was carried out to reduce the dimensionality, the top 30 principal component’s along with a 0.25 resolution were used to cluster and obtain the UMAP embedding of the spots.

### Visium downstream analysis

Clusters were annotated according to pathologists’ assessment, their transcriptional activity and differentially expressed genes. A subset of the single-cell atlas was used to train the *SPOTlight* model, we selected up to 100 cells coming from melanoma cancers for each cell type/state. Selecting cells from one of the cancers types allowed us to reduce dataset-specific noise, which could confound the model. The gene set used to train the model was the union between the marker genes of the cell types along with the top 3000 variable genes. Marker genes for each cell type were obatined with *Seurat’s* function *FindAllMarkers* considering only positive markers and setting the logFC and min.pct to 0 to include all genes. All markers were used to initialize the model basis and unit variance normalization was carried out. Non-smooth nonnegative matrix factorization was the method used to carry out the factorization. Cell types contributing <1% to the spot’s predicted composition were considered fitting noise and were set as 0. We then used *SPOTlight* to map the atlas cell types to the spatial spots.

### Human breast cancer

We used human breast cancer Visium ST data publicly available through the 10x Genomics website (https://support.10xgenomics.com/spatial-gene-expression/datasets/). Two replicates were available and used to confirm the *SPOTlight* predictions. Spatial transcriptomics data run through *spaceranger* 1.0.0 were used (10x Genomics). Further specifications on how the samples were obtained and processed can be found at the 10x Genomics website.

Quality control analysis was carried out looking at the number of unique molecular identifiers (UMIs), genes, and mitochondrial percentage in each spot. No spots were removed from the analysis after inspection of the aforementioned parameters. Data was scaled and normalized using *SCTransform* with default parameters. Principal component analysis was carried out to reduce the dimensionality, the top 30 principal component’s along with a 0.1 resolution were used to cluster and obtain the UMAP embedding of the spots. Clusters were annotated using a pathologist’s annotation to separate tumor from fibrotic. Within tumor, clusters were annotated according to the expression of *ESR1, PGR* and *ERBB2.* We used *SPOTlight* to map the tumor immune cell atlas to the spatial spots. Therefore, a subset was used to train the model, we selected up to 100 cells coming from melanoma cancers for each cell type/state. Selecting cells from one of the cancers types allowed us to reduce dataset-specific noise which could confound the model. The gene set used to train the model was the union between the marker genes of the cell types along with the top 3000 variable genes. Marker genes for each cell type were obatined with Seurat’s function *FindAllMarkers* considering only positive markers and setting the logFC and min.pct to 0 to include all genes. All markers were used to initialize the model basis and unit variance normalization was carried out. Nonsmooth nonnegative matrix factorization was the method used to carry out the factorization. Cell types contributing –1% to the spot’s predicted composition were considered fitting noise and were set as 0. We then used *SPOTlight* to map the atlas cell types to the spatial spots.

### Code versions and availability

All analyses were carried out using R3.6.0 and the data was analyzed using *Seurat* v3.2 (Stuart et al. 2019). Furthermore, *SPOTlight* is developed to run with R versions ≥3.5; docker images with the appropriate environment are available at Docker hub: marcelosua/spotlight_env_rstudio and marcelosua/spotlight_env_r.

