## Supplementary Figures. for "A Single-Cell Tumor Immune Atlas for Precision Oncology"

#### Supplementary Material

**Supplementary Figure legends 1-17**

**Supplementary Figures 1-17**

**Supplementary Figure legends**

**Supplementary Figure 1** | Original annotation of the datasets prior to integration. Azizi et al. 2018 (**A**), Yost et al. 2019 (**B**), Lichun et al. 2019 (**C**), Zhang et al. 2019 (**D, E**), Lee et al. 2020 (**F**), Peng et al. 2019 (**G**), Schelker et al. 2017 (**H**).

**Supplementary Figure 2** | Original annotation of the datasets prior to integration. Lambrechts et al. 2018 (**A**), Lavin et al. 2017 (**B**), Sade-Feldman et al. 2018 (**C**), Li et al. 2019 (**D**), Durante et al. 2020 (**E**), Wu et al. 2020 CRC (**F**), NSCLC (**G**), EA (**H**), RCC (**I**).

**Supplementary Figure 3** | Cell type prediction using a random forest classifier. (**A**) Accuracy and (**B**) kappa statistic using cell type-specific or random signatures as features. (**C**) Confusion matrix displaying the average probability that a cell of a cell type X (rows) is classified as a cell type Y (columns).

**Supplementary Figure 4** | Expression of selected markers across all cell types.

**Supplementary Figure 5** | Immune cell states correlate in composition frequencies and enable to cluster patients into immune subtypes. (**A**) Correlation matrix of the 25 immune cell states across patients. Co-occurring and mutually exclusive relationships are quantified by Pearson's correlation (positive: red; negative: blue). (**B-D**) Dotplots depicting the percentage of cells that express and the expression level of cluster marker/exhaustion genes. Plots display the top 10 marker genes for each immune cluster (C1-6, B), the expression of exhaustion markers in CD8 T-cells split by immune cluster (C1-6, C) and the expression of exhaustion marker in all cell states in cluster 5 (C5, D).

**Supplementary Figure 6** | Random Forest top 15 most significant variables.

**Supplementary Figure 7** | SPOTlight scheme. Step-by-step illustration of SPOTlight's algorithm. At the beginning of this process we have a count matrix, V, for scRNAseq data and a set of marker genes for the identified cell types. First, we use prior information to initialize the basis and coefficient matrices, W and H respectively. We assume the number of topics, k, to be equal to the number of cell types in the dataset. Each topic is then associated with a cell type; columns in W are initialized with marker genes for the associated cell type with that topic, while rows in H are initialized with the membership of each cell to its associated topic. Second, we proceed with the matrix factorization from which we obtain gene distributions for each topic in W, and topic profiles for each cell in H. Third, we use W to map the ST data, V', by means of non-negative least squares (NNLS) to obtain H'. Columns in H' represent the topic profile for each spot. Fourth, from the H matrix obtained from the scRNAseq data we consolidate all the cells from the same cell type to obtain cell type-specific topic profiles. Lastly, we use NNLS to find which combination of cell type-specific topics resembles each spot's topic profile.

**Supplementary Figure 8** | Sample 161429 (SCC) predicted cell type/state proportion within each spot showing spatially differential immune patterns.

**Supplementary Figure 9** | Sample 161429 (SCC) box plots of the cell types/states of interest presenting differential proportions of cell types found between regions. Differences between tumor sub-regions suggest a differential tumor immune microenvironment between heterogeneous tumor regions.

**Supplementary Figure 10** | Tumor immune cell atlas mapping on Visium oropharyngeal cancer, sample ID 161430. (**A**) Cell type/state specific topic profiles showing a high topic – cell type/state specificity. (**B**) Tissue stratification and labeling according to unsupervised clustering (**C**) Number of unique molecular identifiers (UMI) recovered from each Visium spot suggesting which areas might be more transcriptionally active. (**D**) Pie chart representation of each spot showing the immune cells that mapped to it. Shown are immune cell types present in <90% of the spots so as to present those that are variable between regions. (**E**) UMAP embedding of the spots presenting the cell cycle phase each spot is in, the clustering, and the UMI counts. (**F**) Box plots of the cell types of interest representing differential proportions of cell types found between regions. Differences between tumor sub-regions are present suggesting a differential tumor immune microenvironment between heterogeneous tumor regions. (**G**) Location and proportion of cell types of interest on the tissue. (**H**)

Clustered correlation matrix between the predicted cell type/state proportions showing clear correlating and anti-correlating cell types. Excluded are cell types/states not predicted to be present on the tissue. (**I**) scatter plots of correlating cell type pairs of interest.

**Supplementary Figure 11** | Sample 161430 predicted cell type/state proportion within each spot showing spatially differential immune patterns.

**Supplementary Figure 12** | Sample 161430 box plots of the cell types/states of interest presenting differential proportions of cell types found between regions. Differences between tumor sub-regions suggest a differential tumor immune microenvironment between heterogeneous tumor regions.

**Supplementary Figure 13** | TICA mapping on Visium ductal carcinoma breast tissue spatial slice 2 data. (**A**) Estrogen receptor gene (ESR1) expression on the tissue slide. Clear regionality of the expression is seen indicating ER+ regions. (**B**) HER2 receptor gene (ERBB2) expression on the tissue slide. Clear regionality of the expression is seen indicating HER2+ regions. (**C**) Tissue stratification and labeling according to unsupervised clustering and ER and HER2 expression. (**D**) Pie chart representation of each spot showing the immune cells that mapped to it. Shown are immune cell types present in <90% of the spots so as to present those that are variable between regions. (**E**) Box plots of the cell types of interest representing differential proportions of cell types found between regions. Differences between tumor sub-regions are present suggesting a differential tumor immune microenvironment between the heterogeneous tumor regions. (**F**) Location and proportion of cell types of interest on the tissue. (**G**) Clustered correlation matrix between the predicted cell-type proportions showing clear correlating and anti-correlating cell types. Excluded are cell types/states not predicted to be present on the tissue. (**H**) scatter plots of correlating cell type pairs of interest.

**Supplementary Figure 14** | Predicted cell type/state proportion within each spot showing spatially differential immune patterns for the breast ductal carcinoma slice 1.

**Supplementary Figure 15** | Predicted cell type/state proportion within each spot showing spatially differential immune patterns for the breast ductal carcinoma sequential slice 2 replicate.

**Supplementary Figure 16** | Box plots of the cell types/states of interest presenting differential proportions of cell types found between regions for the breast ductal carcinoma slice 1. Differences between tumor sub-regions suggest a differential tumor immune microenvironment between heterogeneous tumor regions.

**Supplementary Figure 17** | Box plots of the cell types/states of interest presenting differential proportions of cell types found between regions for the breast ductal carcinoma sequential slice 2 replicate. Differences between tumor sub-regions suggest a differential tumor immune microenvironment between heterogeneous tumor regions.

Supplementary Figure 1

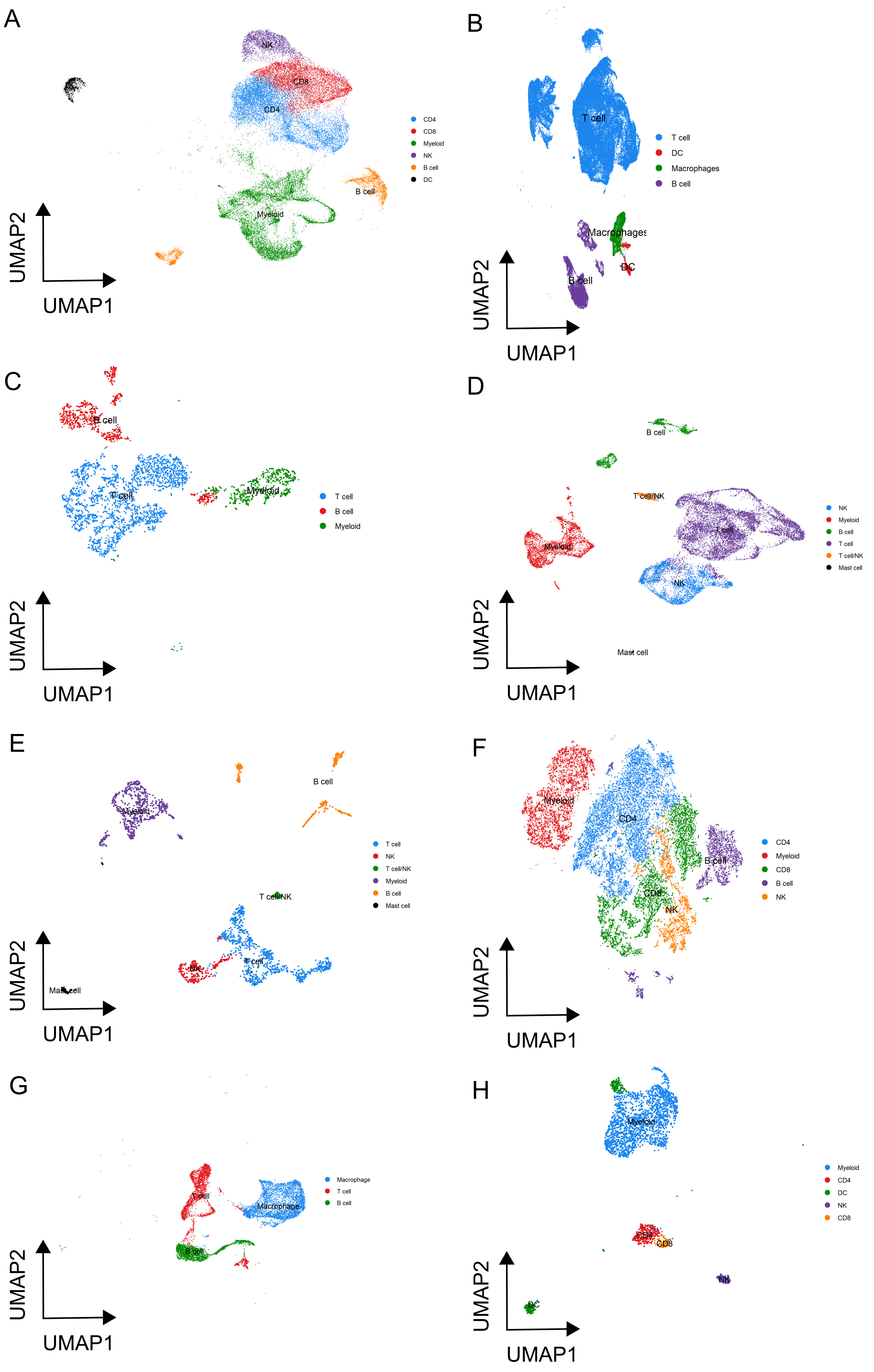

Supplementary Figure 2

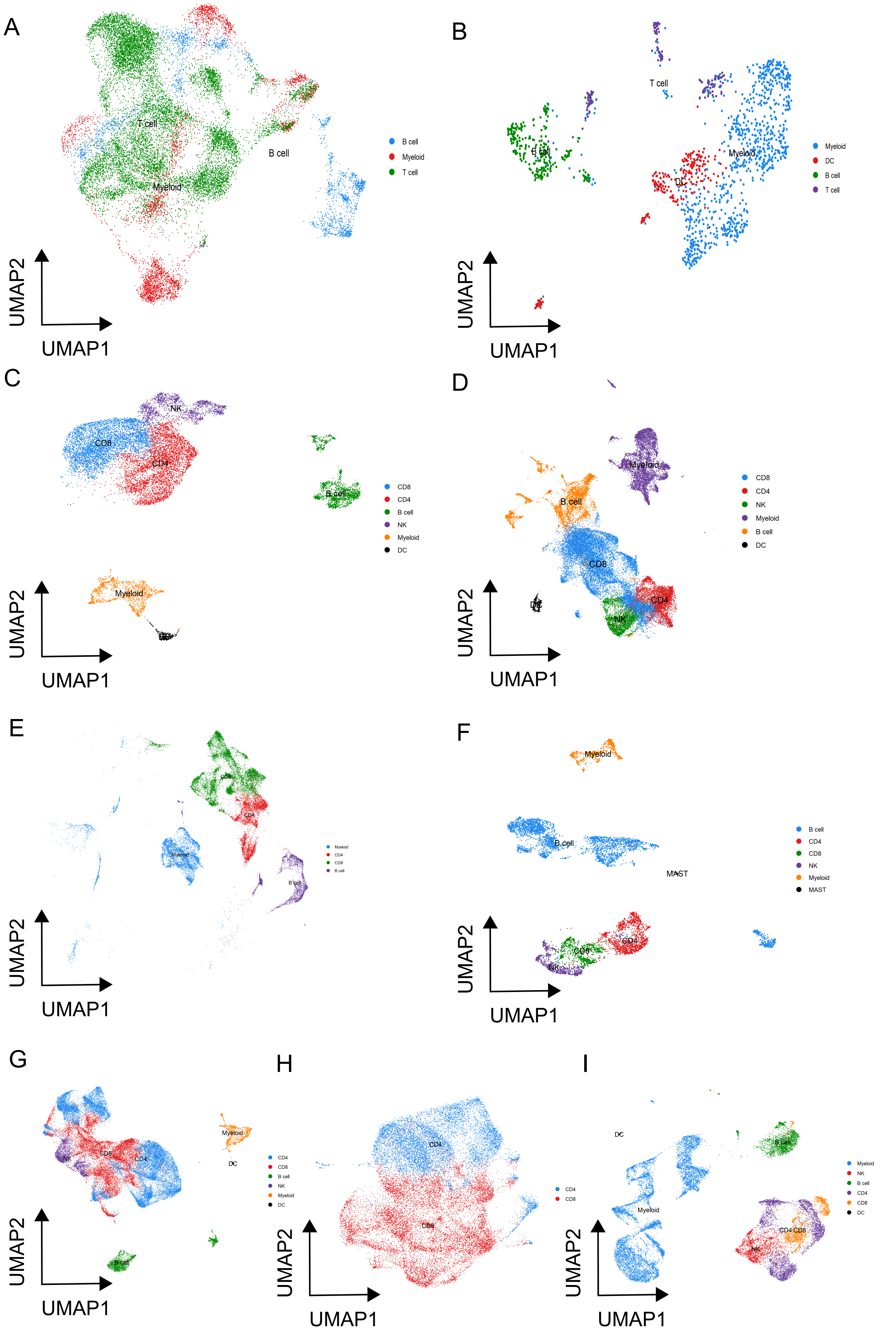

Supplementary Figure 3

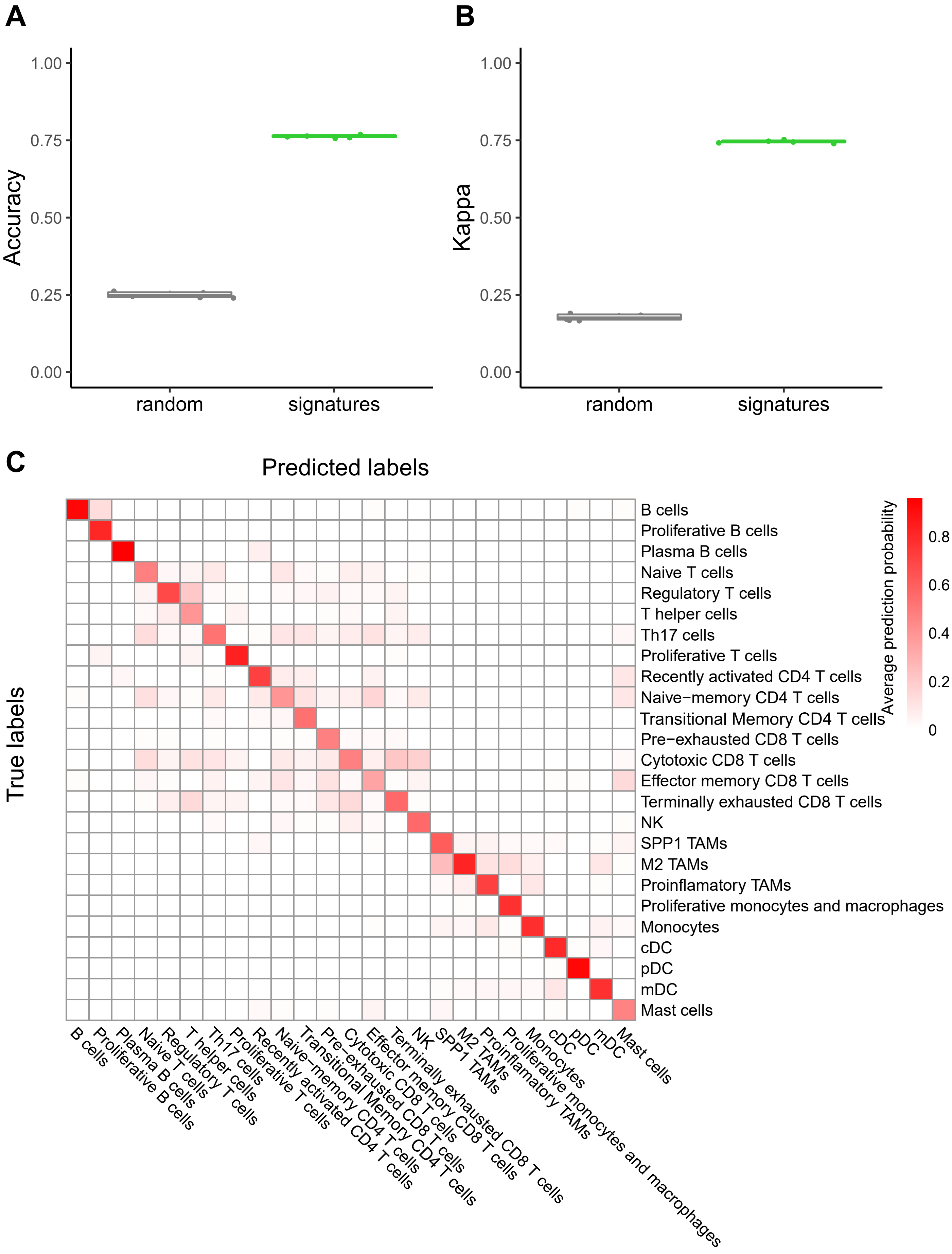

Supplementary Figure 4

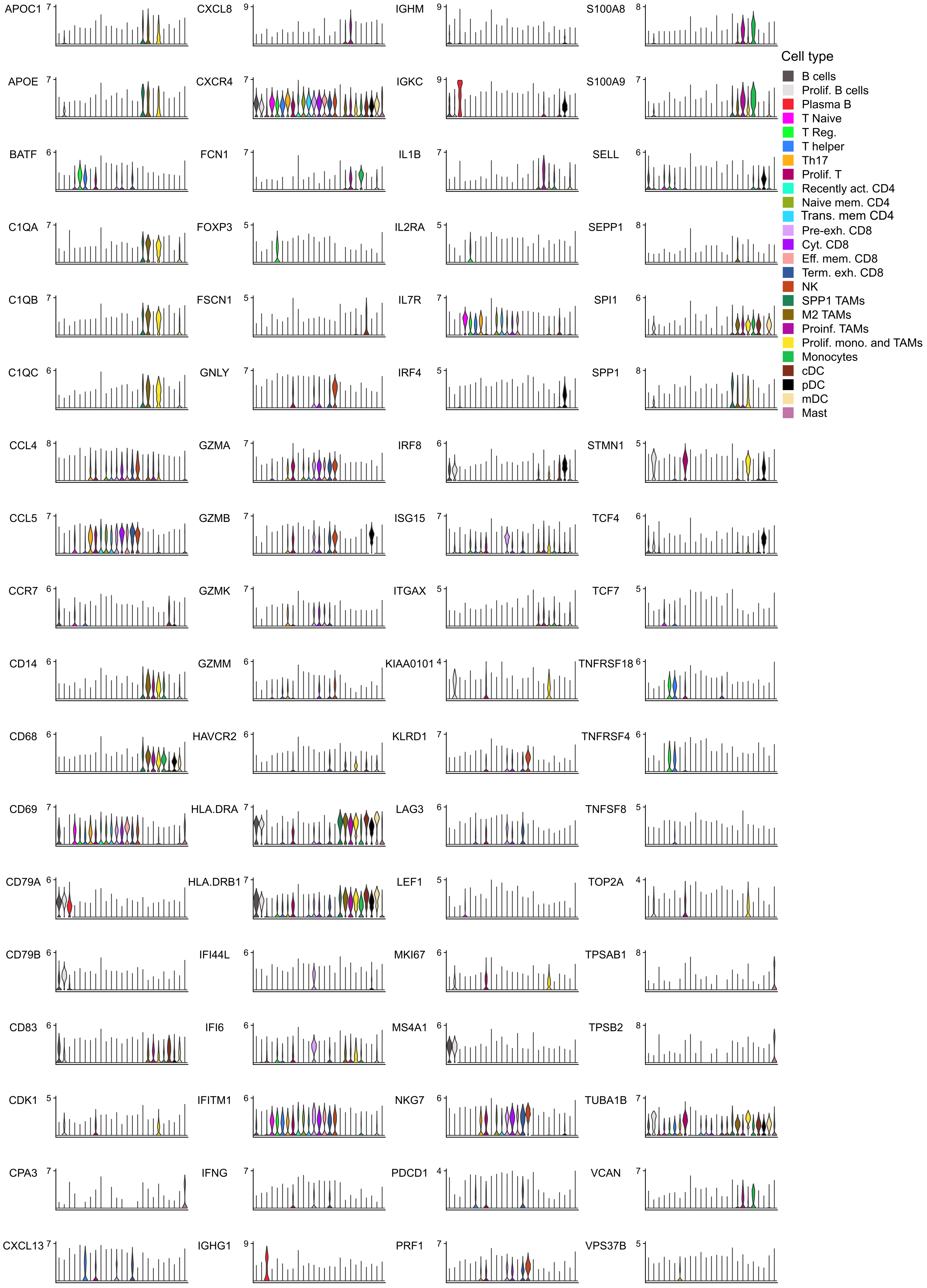

### Supplementary Figure 5

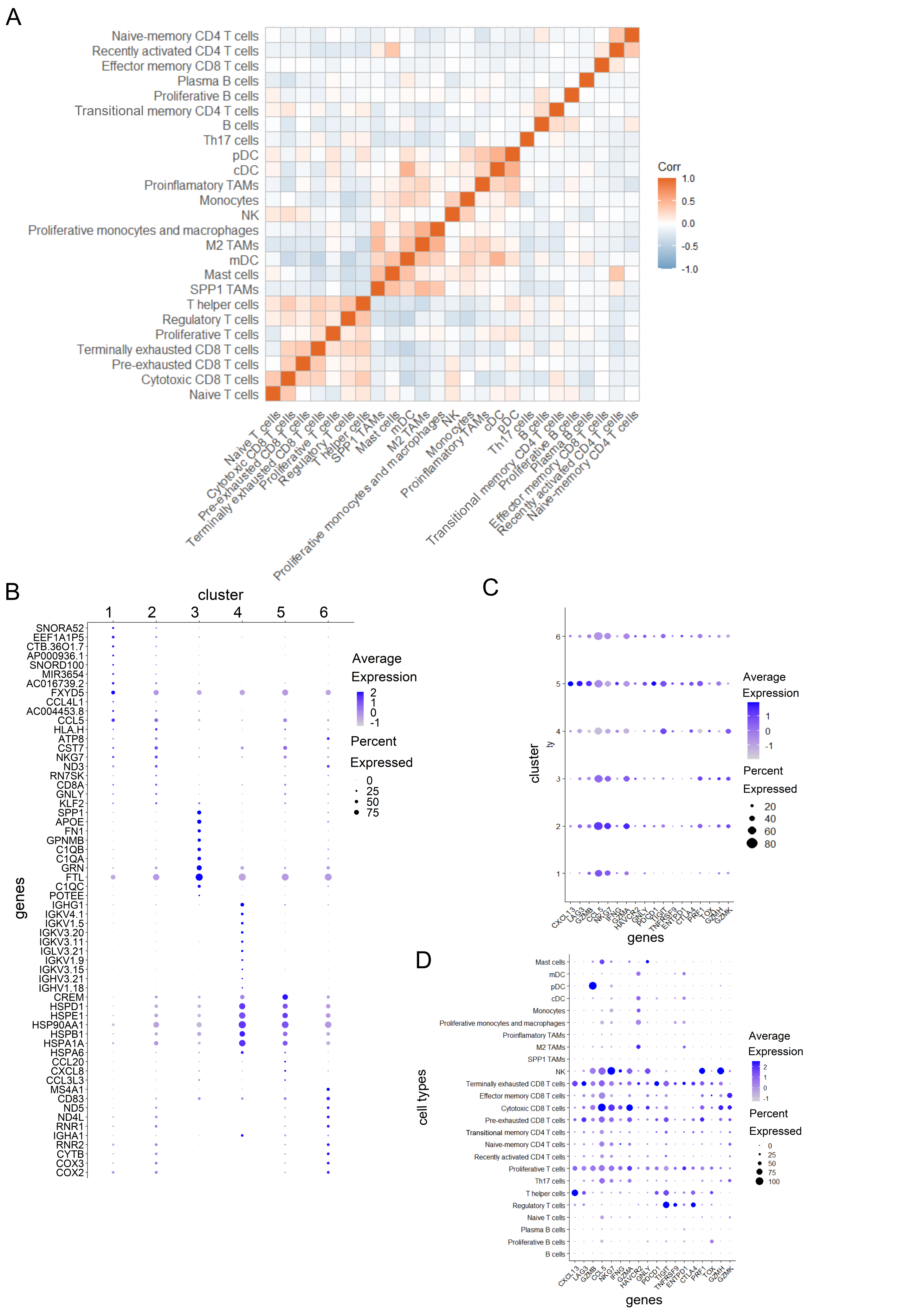

A

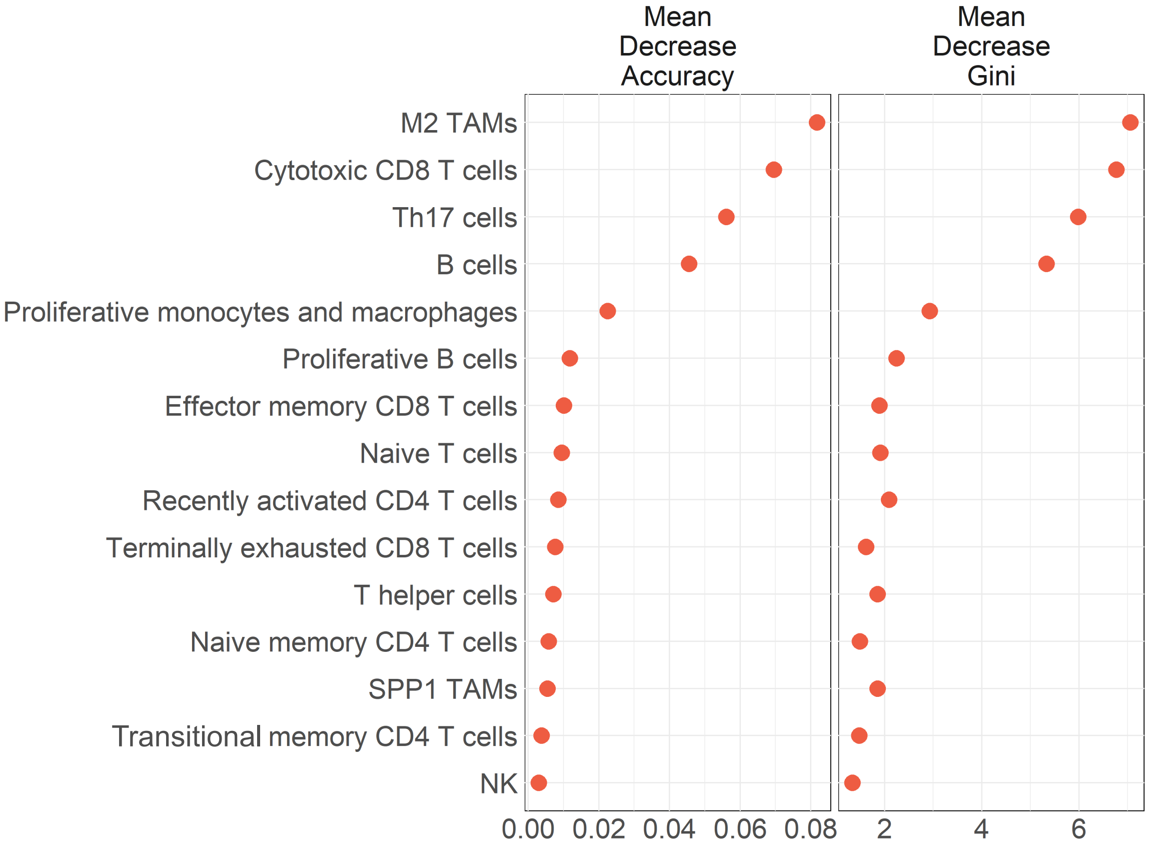
Mean Decrease Accuracy
Mean Decrease Gini
M2 TAMs
Cytotoxic CD8 T cells
Th17 cells
B cells
Proliferative monocytes and macrophages
Proliferative B cells
Effector memory CD8 T cells
Naive T cells
Recently activated CD4 T cells
Terminally exhausted CD8 T cells
T helper cells
Naive memory CD4 T cells
SPP1 TAMs
Transitional memory CD4 T cells
NK
0.00 0.02 0.04 0.06 0.08
2 4 6

B

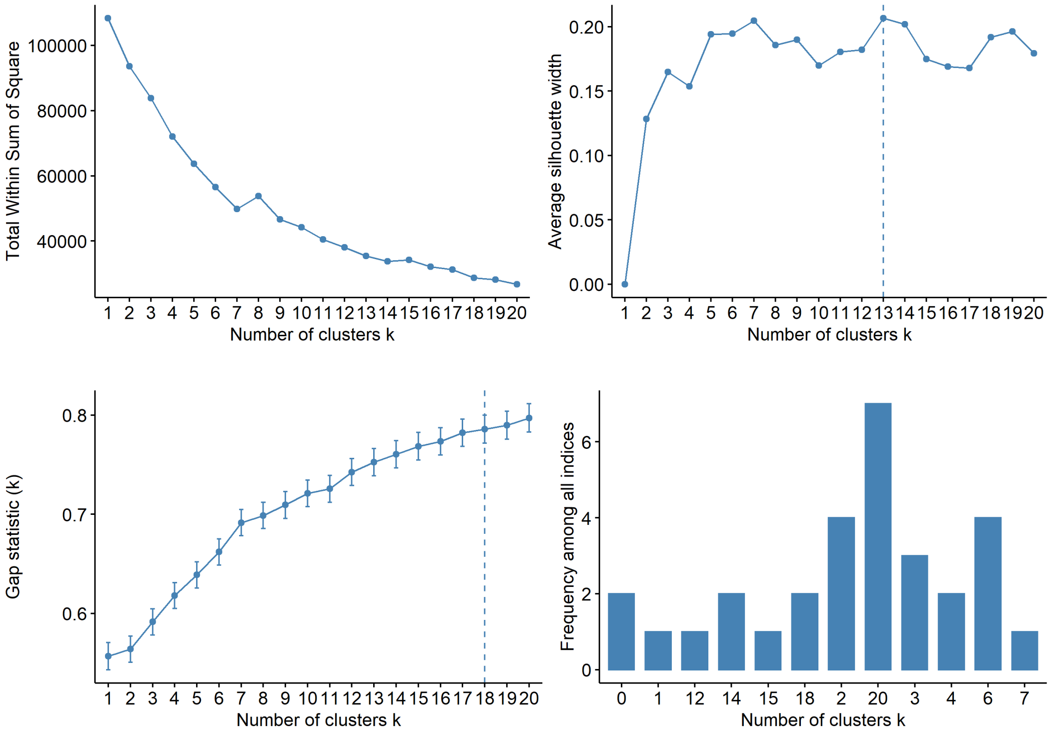
Total Within Sum of Square
Number of clusters k
Average silhouette width
Number of clusters k
Gap statistic (k)
Number of clusters k
Frequency among all indices
Number of clusters k

Supplementary Figure 7

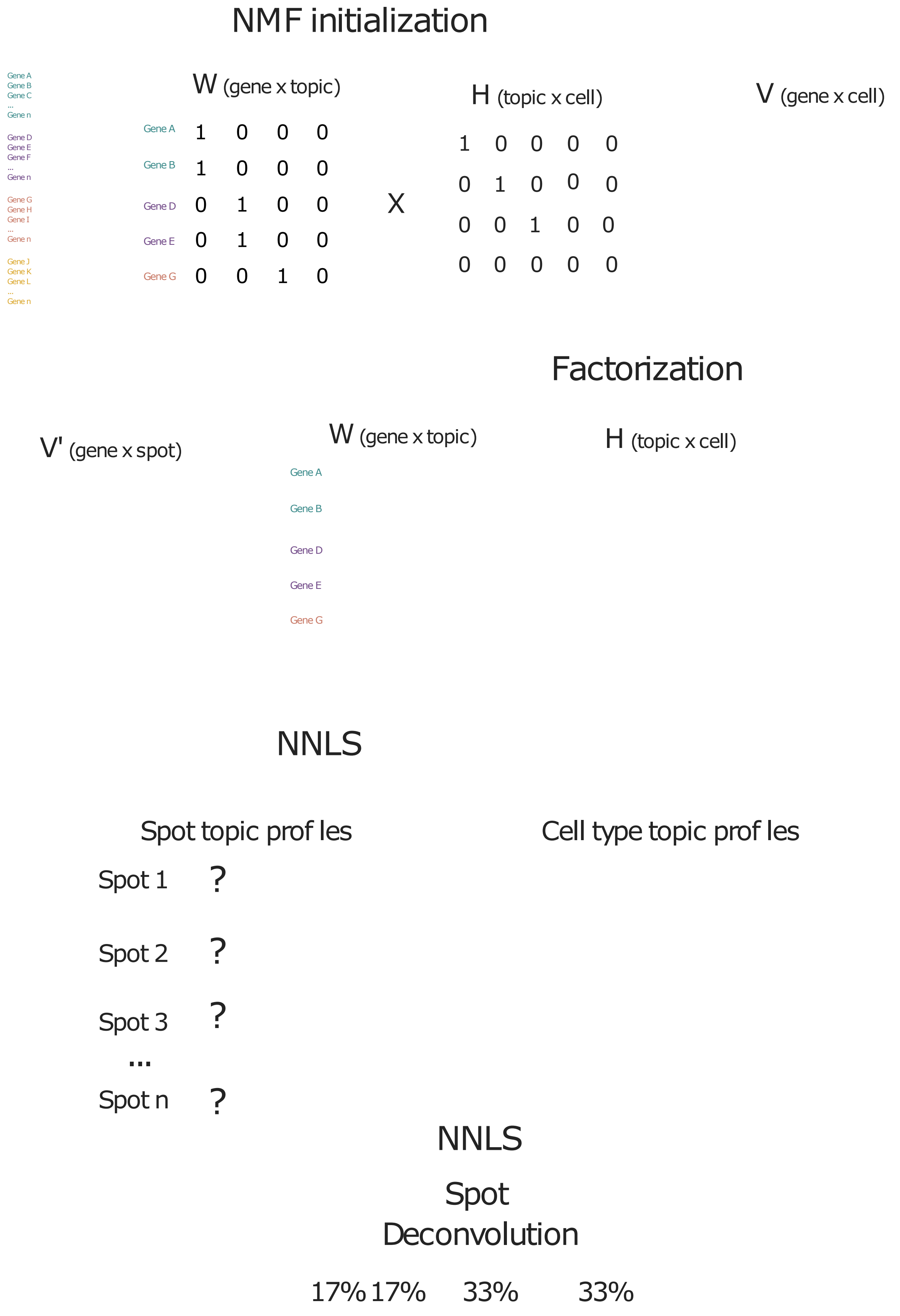

Supplementary Figure 8

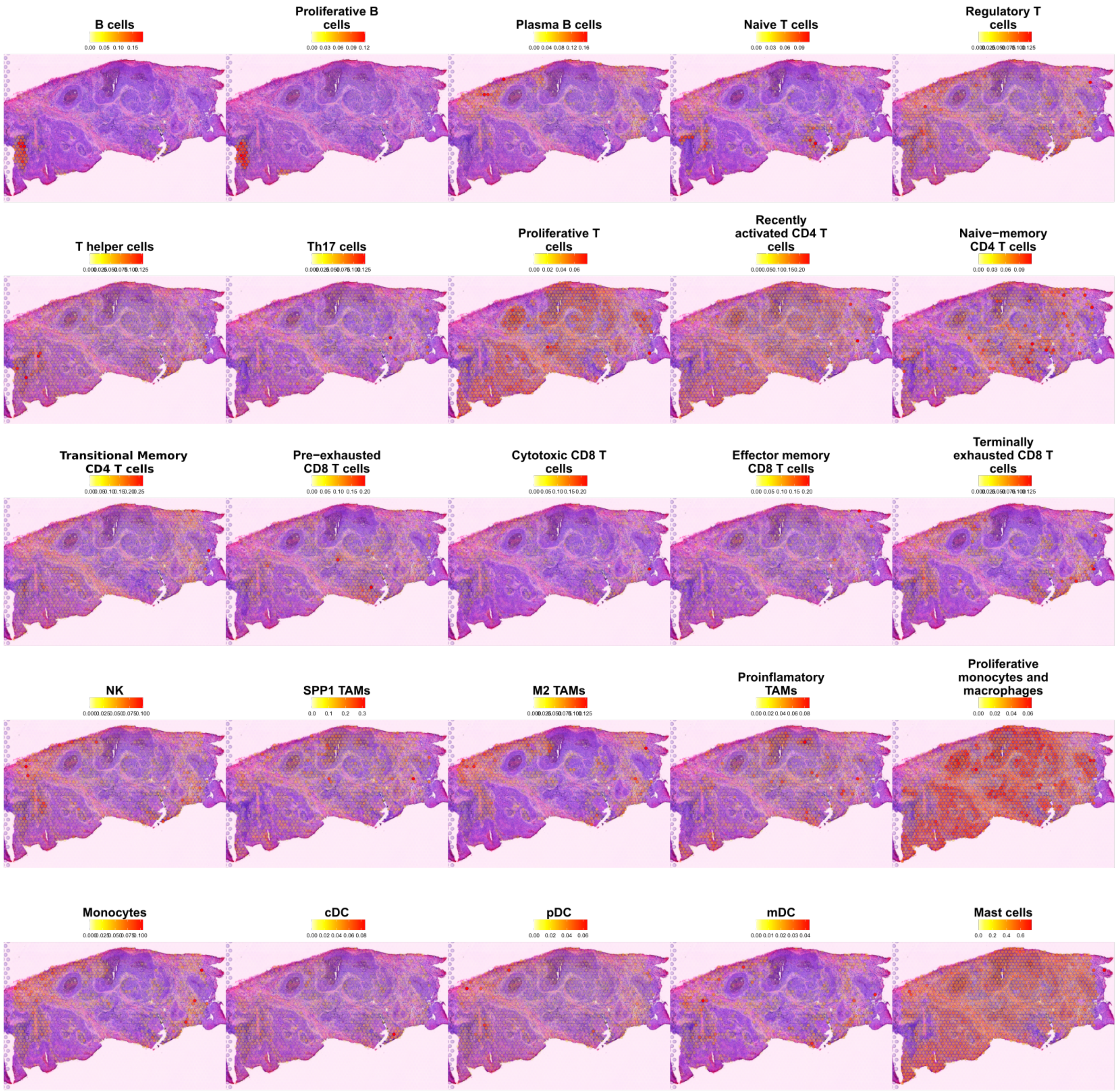

### Supplementary Figure 9

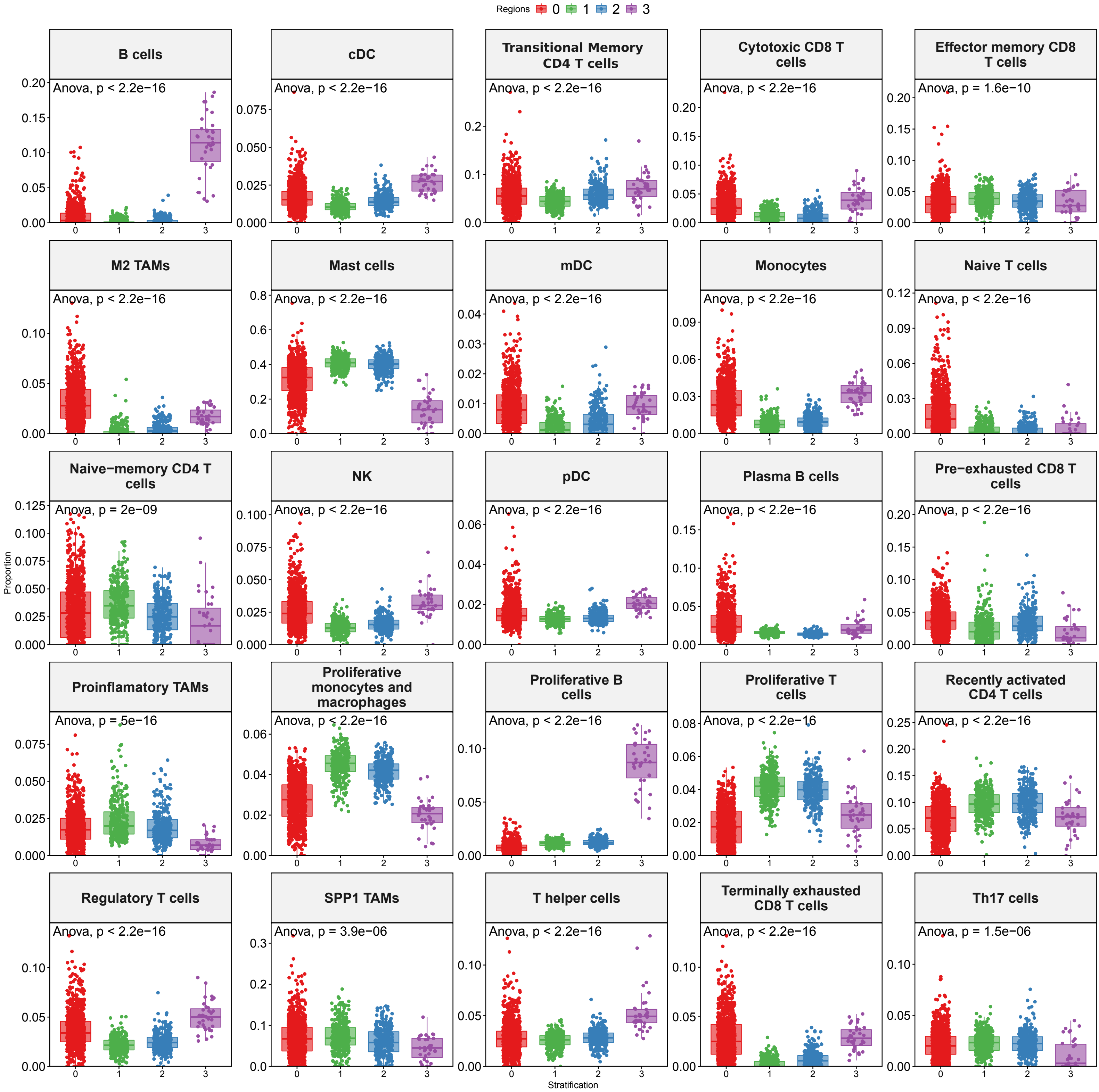

Supplementary Figure 10

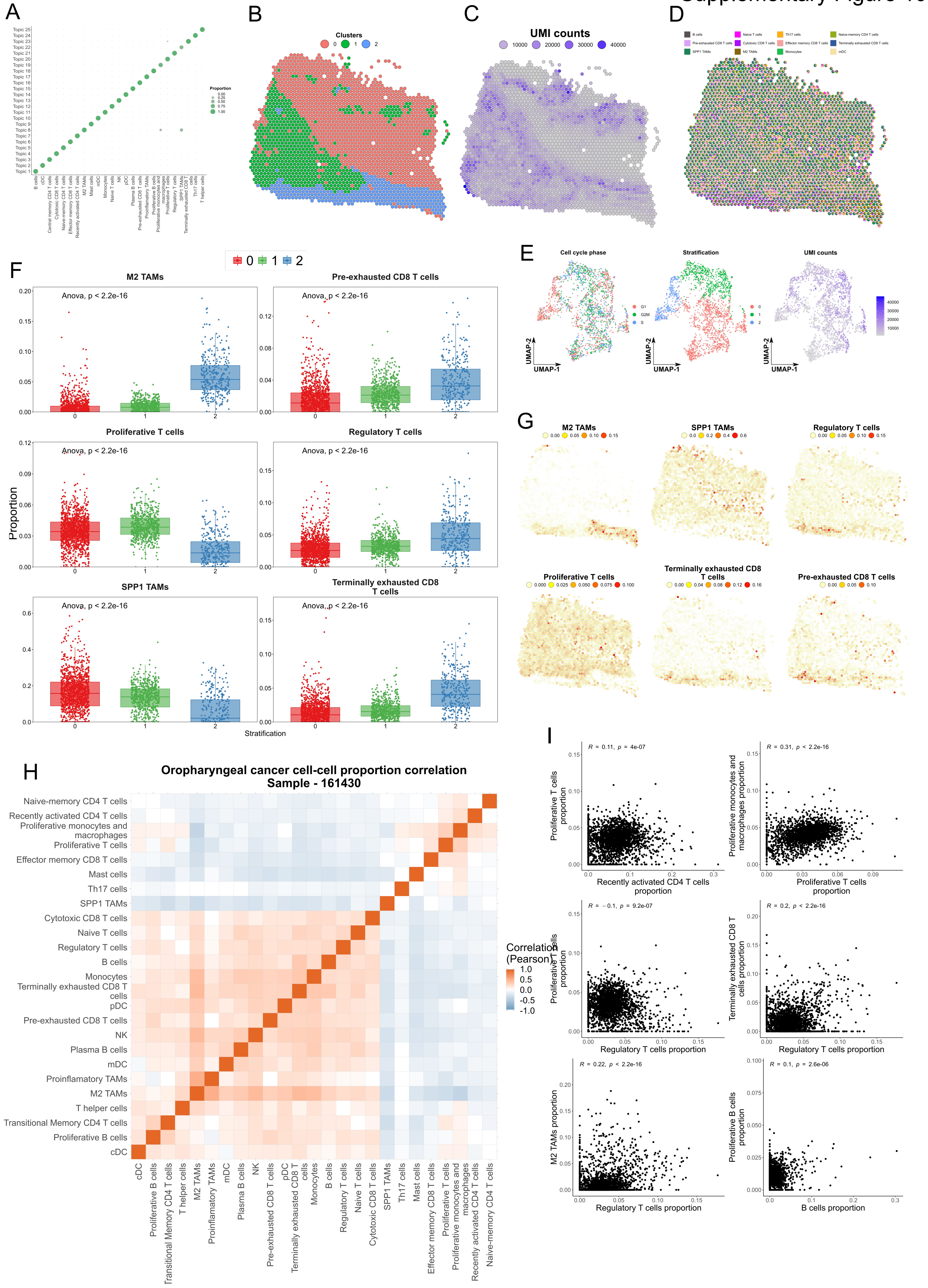

Supplementary Figure 11

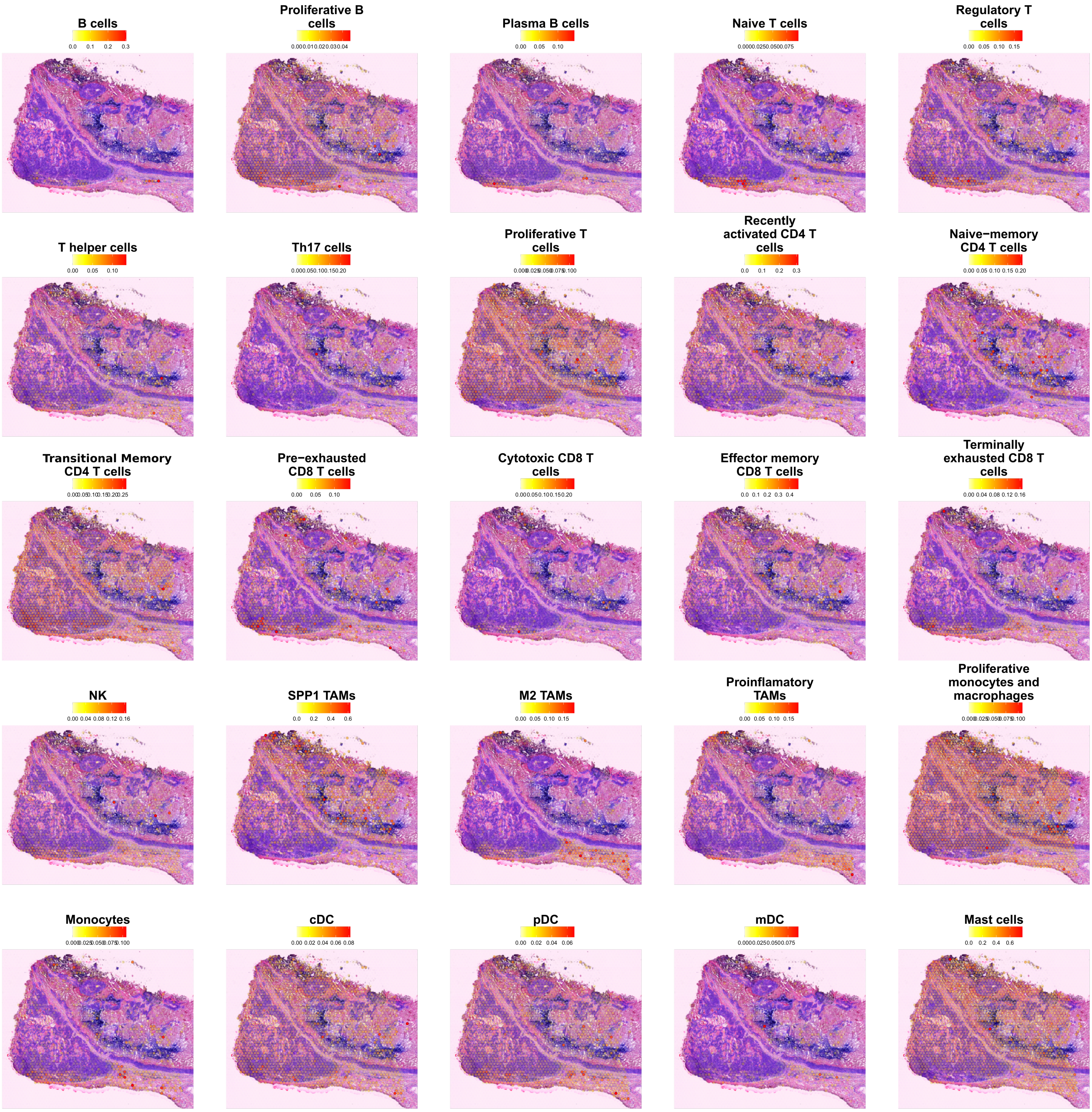

Supplementary Figure 12

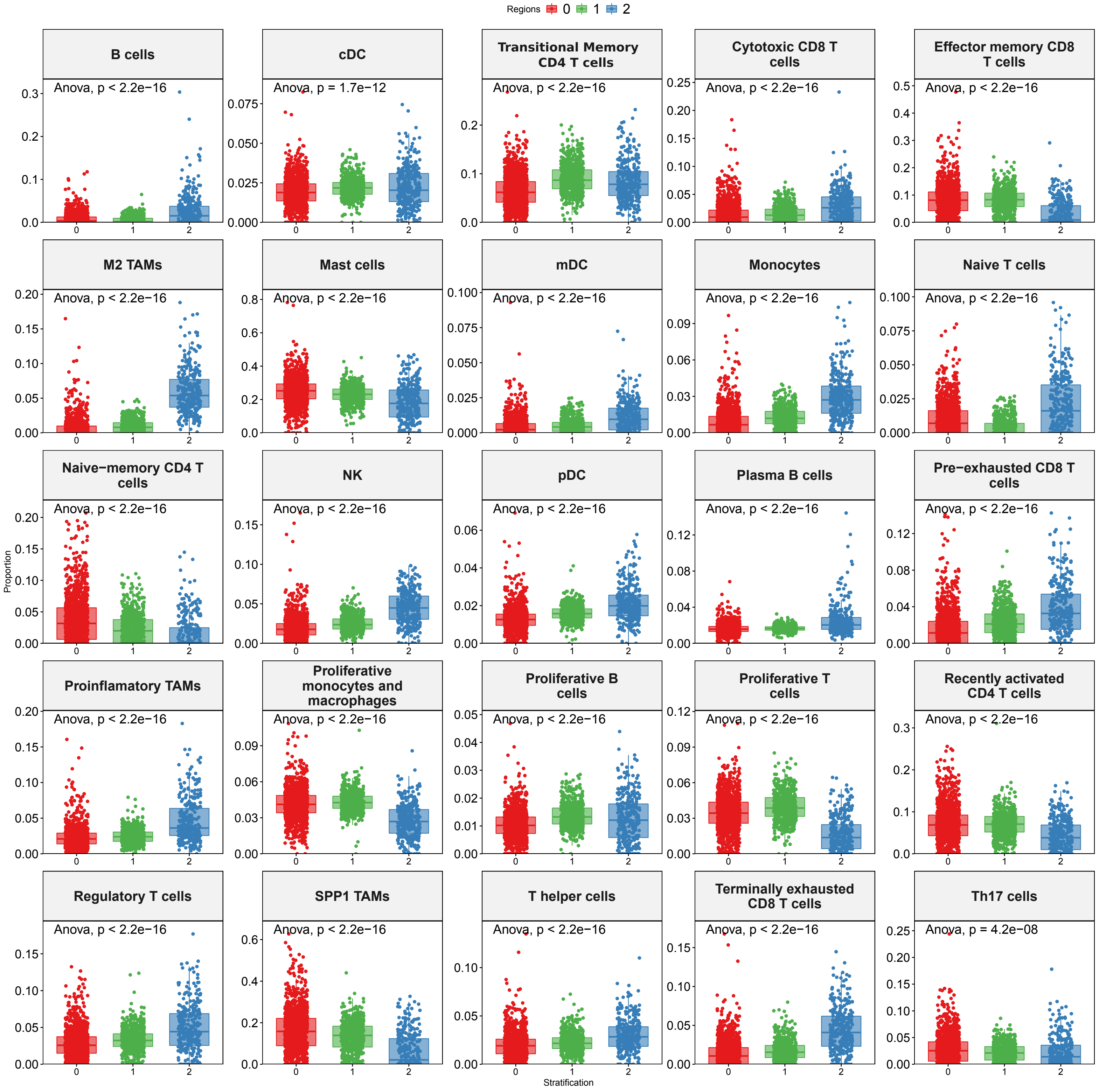

### Supplementary Figure 13

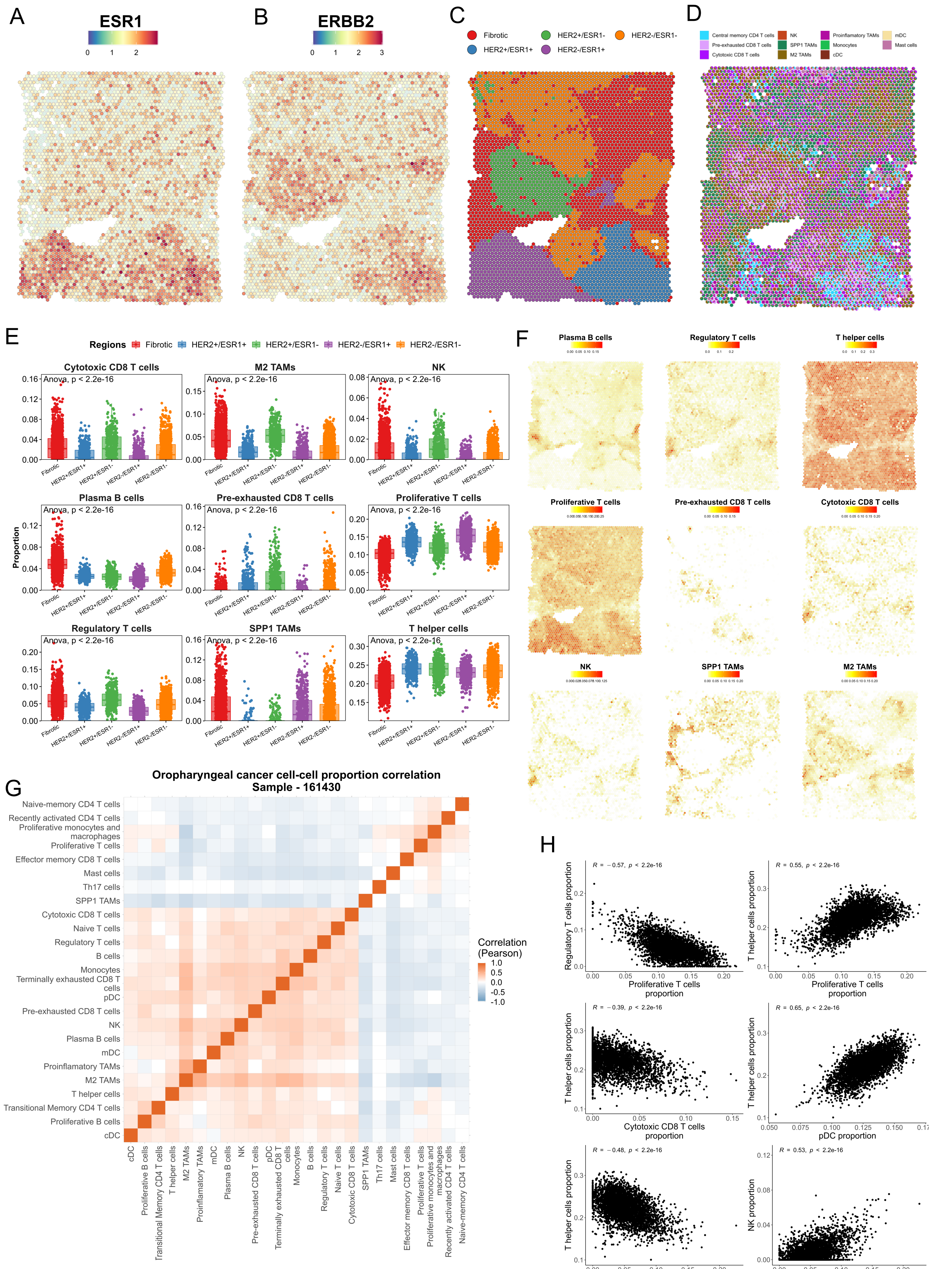

Supplementary Figure 14

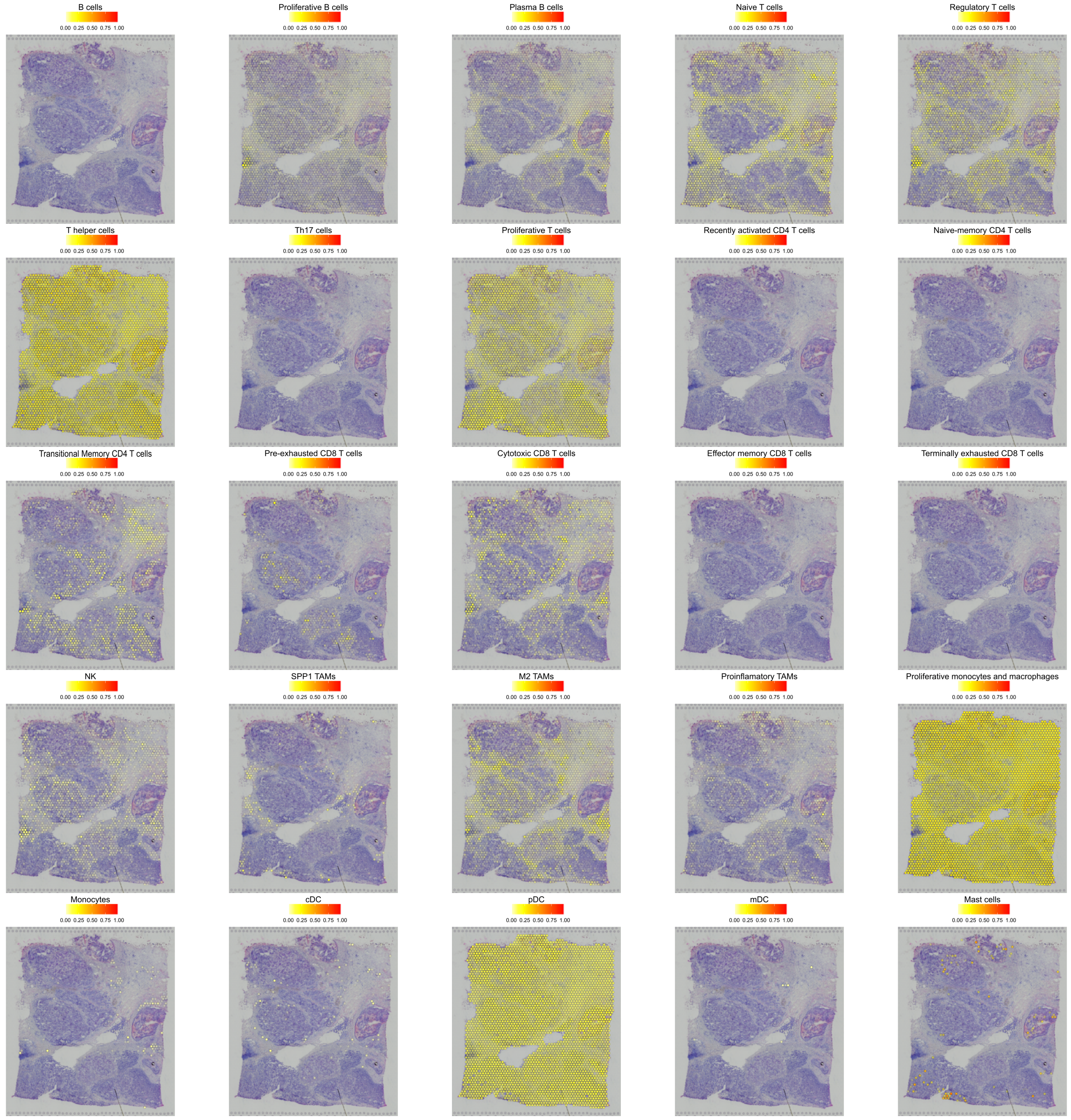

### Supplementary Figure 15

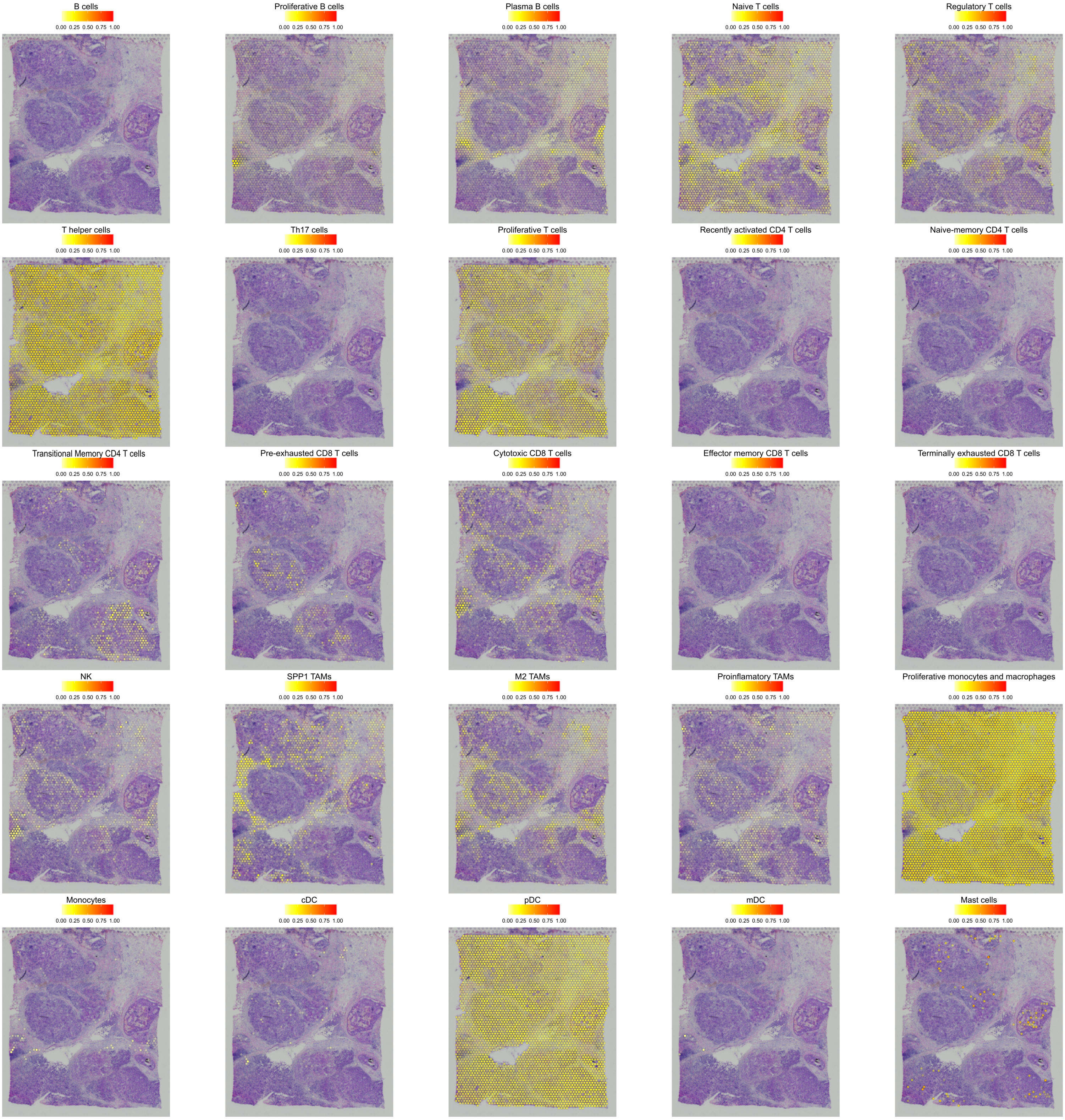

### Supplementary Figure 16

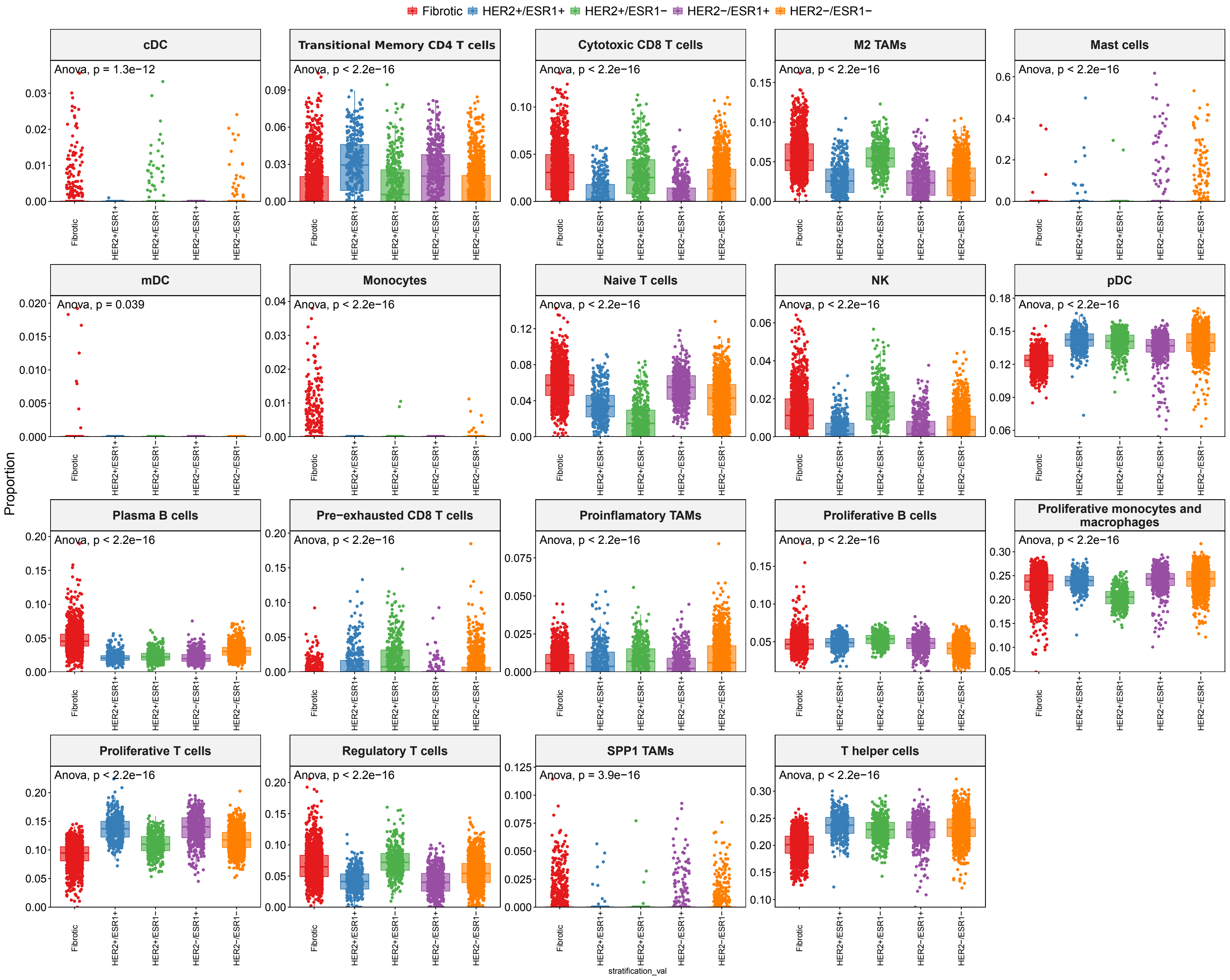

### Supplementary Figure 17

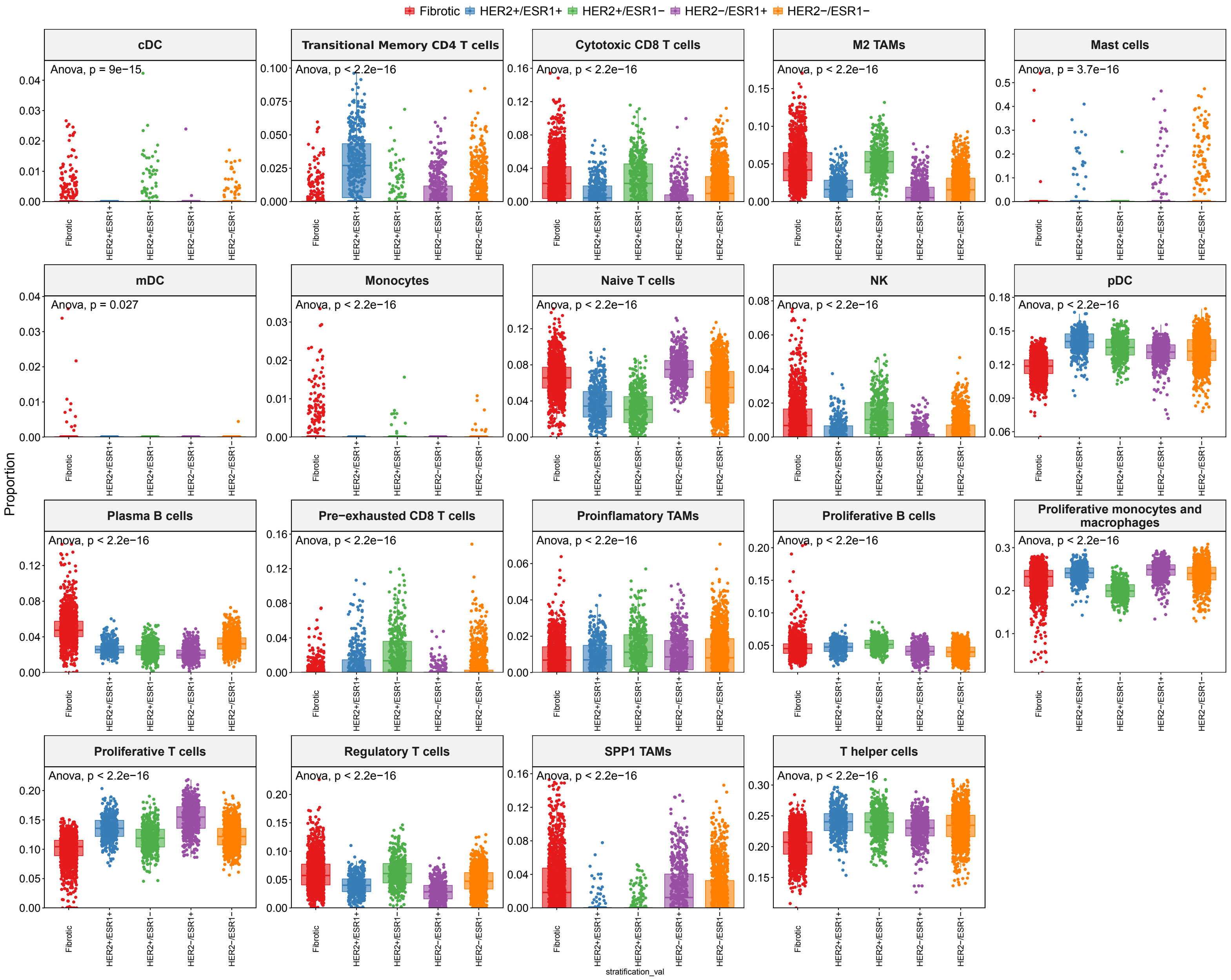
