## Supplementary_Table1 for "A Single-Cell Tumor Immune Atlas for Precision Oncology"

| **Paper** | **Cancer type** | **# patients** | **# total cells** | **# immune cells** | **Sequencing technology** |
| --- | --- | --- | --- | --- | --- |
| Azizi et al. 2018 | breast carcinoma (BC) | 8 | 46,642 | 28,918 | inDrop |
| Yost et al. 2019 | basal cell carcinoma (BCC) and squamous cell carcinoma (SCC) | 15 | 79,046 | 59,792 | 10x |
| Lichun et al. 2019 | intrahepatic cholangiocarcinoma (ICC) and hepatocellular carcinoma (HCC) | 19 | 9,536 | 4,331 | 10x |
| Zhang et al. 2019 | intrahepatic cholangiocarcinoma (ICC) and hepatocellular carcinoma (HCC) | 16 | 20,393 | 17,242 | 10x and Smart-Seq2 |
| Lee et al. 2020 | colorectal cancer (CRC) | 43 | 63,689 | 31,367 | 10x |
| Peng et al. 2019 | pancreatic ductal adenocarcinoma (PDAC) | 24 | 41,986 | 10,623 | 10x |
| Schelker et al. 2017 | ovarian cancer (OC) | 4 | 2,765 | 2,432 | inDrop |
| Lambrechts et al. 2018 | non-small-cell lung cancer (NSCLC) | 5 | 24,908 | 17,077 | 10x |
| Lavin et al. 2017 | non-small-cell lung cancer (NSCLC) | 28 | 1,519 | 1,460 | MARS-Seq |
| Sade-Feldman et al. 2018 | melanoma | 29 | 16,188 | 14,991 | Smart-Seq2 |
| Li et al. 2019 | melanoma | 22 | 44,403 | 37,398 | MARS-Seq |
| Durante et al. 2020 | uveal melanoma | 11 | 86,407 | 8,513 | 10x |
| Wu et al. 2020 | colorectal cancer (CRC) | 2 | 7,138 | 6,909 | 10x |
|  | non-small-cell lung cancer (NSCLC) | 6 | 39,323 | 35,107 |  |
|  | endometrial adenocarcinoma (EA) | 6 | 22,706 | 22,611 |  |
|  | renal cell carcinoma (RCC) | 3 | 19,612 | 18,340 |  |
