## Supplementary_Table2 for "A Single-Cell Tumor Immune Atlas for Precision Oncology"

| **cell type** | **markers** |
| --- | --- |
| B cells | MS4A1, CD79A, CD83, CD79B, CD37, CD19 |
| Proliferative B cells | CD79B, MS4A1, CD79A, STMN1, KIAA0101, TUBA1B, CD19 |
| Plasma B cells | IGKC, IGHG1, IGHM, TNFRSF17, SDC1, CD38 |
| Naive T cells | IL7R, TCF7, CCR7, LEF1, SELL |
| Regulatory T cells | FOXP3, TNFRSF18, IL2RA, TIGIT, CTLA4, IKZF2 |
| T helper cells | CXCL13, TNFRSF4, TNFRSF18, BATF, TIGIT, SOX4, TNFRSF25, CTLA4, RORA, XCL1, TNFSF8, PPIA, STAT5A, TOX, PDCD1 |
| Th17 cells | TNFSF8, IL7R, CXCR4, VPS37B, GZMK, STAT4, TGFB1, XCL1, XCL2, CCR7, CRTAM |
| Proliferative T cells | STMN1, MKI67, CDK1 |
| Recently activated CD4 T cells | CCL4, IFITM1, CD69, PRF1, BCL3, IL7R, IFITM3, TCF7, CD81, CXCR4, GZMK, GZMM, IFITM2 |
| Naive memory CD4 T cells | IL7R, TCF7, CCL5, IFITM1 |
| Transitional Memory CD4 T cells | CD2, CD28, CD44, CD6, CD69, CD96, CTLA4, IL6ST, IL7R, FLNA, SESN3 |
| Pre-exhausted CD8 T cells | ISG15, IFI44L, IFI6, IFIT3, IFIT1, IFI44, IFI35, IRF7, IFIT2, LAG3, IFITM1, IFI16, IFI27, IFNG, GZMB, GZMK, PRF1, HAVCR2, IFIH1, GZMA, IRF9, CXCL13, GZMH, IFIT5, PDCD1 |
| Cytotoxic CD8 T cells | CCL5, GZMA, GZMK, NKG7, GNLY, GZMH, GZMM, PRF1, CXCR3, GZMB, CCL4, IFNG |
| Effector memory CD8 T cells | GZMM, IFITM1, GZMK, IFNG, CCL5 |
| Terminally exhausted CD8 T cells | CXCL13, LAG3, GZMB, CCL5, NKG7, IFNG, GZMA, HAVCR2, GNLY, PDCD1, TIGIT, TNFRSF9, ENTPD1, CTLA4, PRF1, TOX, GZMH, GZMK |
| NK | PRF1, GNLY, KLRD1, KLRF1, GZMH, GZMB, KLRB1, GZMA, GZMM, CD160, CD244, KLRC1, NCR1 |
| SPP1 TAMs | SPP1, APOE, SEPP1, MMP9, CD163 |
| M2 TAMs | C1QB, APOE, C1QA, C1QC, APOC1, SEPP1, SPP1, CD163 |
| Proinflamatory TAMs | CXCL8, IL1B, S100A9, S100A8, CCL2, IL8, CD68, IL6, IL1A |
| Proliferative monocytes and macrophages | C1QB, C1QC, SPP1, C1QA, TUBA1B, STMN1, APOE, APOC1, MKI67, CD14, KIAA0101, TOP2A, CD68, SPI1, BIRC5, CSF1R, PCNA, FCER1A, IL1B, CD1C |
| Monocytes | S100A8, S100A9, FCN1, VCAN, AIF1, SPI1, CD14, APOBEC3A, CSF1R, ASAH1 |
| cDC | FSCN1, HLA.DRA, HLA.DRB1, SPI1, CLEC9A, XCR1, ITGAX |
| pDC | TCF4, IRF8, IRF4, BCL11A, SPIB, CLEC4C, RUNX2 |
| mDC | HLA.DRA, HLA.DRB1, SPI1, CD68, CD83, ITGAX, CD1D |
| Mast cells | TPSB2, TPSAB1, CPA3, GATA2, KIT, MS4A2 |
